# Carbon source diversity shapes bacterial interspecies interactions

**DOI:** 10.1101/2025.03.30.646244

**Authors:** Hiroki Ono, Saburo Tsuru, Chikara Furusawa

## Abstract

Bacterial communities exhibit various classes of interspecies interactions, ranging from cooperative to competitive. As these interaction classes play a crucial role in determining characteristics of bacterial communities, including species composition and community stability, understanding of the mechanisms that shape them is warranted. While several studies have suggested that cooperative interactions are rare, a study focused on single-carbon-source environments reported them to be relatively common. This discrepancy highlights the potential role of carbon source diversity in shaping interaction classes, although the quantitative relationship remains unclear. To elucidate this relationship, we examined 896 interspecies interactions among 28 synthetic bacterial pairs, isolated from various environments, under 32 conditions with varying levels of carbon source diversity. As a result, we frequently observed cooperative interactions in single-carbon-source environments, with the interactions shifting to competitive as the carbon source diversity increased. Further analyses suggested that this shift was driven by intensified resource competition in more complex media. Our findings provide new insights into how environmental factors, particularly carbon source diversity, shape bacterial interactions in synthetic communities, offering potential strategies for manipulating bacterial communities in ecological and biotechnological contexts.

## Introduction

Many bacteria coexist with multiple species, forming complex communities where they interact either indirectly through their growth environment [1, 2] or directly via physical contact [3, 4]. These interspecies interactions include various classes [5–7], ranging from competition, in which both species inhibit growth of each other, to mutualism, in which they promote growth for both species. The interaction classes affect species composition and community stability [8–14], which in turn plays a key role in essential functions of natural communities, such as global nutrient cycling [15] and food digestion [16], as well as applications of synthetic communities, including material production [17], bioremediation [18], and bacteriotherapy [19]. Therefore, it is crucial to quantitatively understand how different interaction classes emerge under specific conditions for the effective utilisation of both natural and synthetic communities [20, 21].

Toward understanding the principles governing interaction classes, previous studies primarily focused on combinations of bacterial pairs and growth environments. Several studies simulated bacterial interactions under various growth environments, suggesting that cooperative interactions—such as mutualism (bidirectional growth facilitation) and commensalism (unidirectional growth facilitation)—can form to some extent via their metabolites produced in response to the growth environments [22–25]. Furthermore, the Black Queen Hypothesis [26] suggests that cooperative interactions mediated by such metabolites may be ubiquitous in bacterial communities.

Many studies that experimentally assessed bacterial interactions, however, reported that cooperative interactions were rare [27–31]. In these studies, bacterial interactions were examined using synthetic pairs of two isolated species. Notably, each of these studies examined a single natural medium containing multiple carbon sources and reported that competitive interactions—such as competition (bidirectional growth inhibition) and amensalism (unidirectional growth inhibition)—occurred predominantly. These results made it seem that cooperative interactions via secreted metabolites, which were theoretically predicted to occur frequently, were rare [32].

In contrast to these previous studies, Kehe et al. [33] investigated synthetic bacterial pairs under well-defined media containing only a single carbon source selected from 33 different organic compounds. Their findings revealed that cooperative interactions were more common in single-carbon-source environments, contrasting with studies conducted in multi-carbon-source environments [27–31]. The discrepancies in these results suggest that the growth environment, especially carbon source diversity, can affect bacterial interaction classes. However, few studies have systematically examined bacterial interactions across different environments, particularly those with varying carbon source diversity while controlling for other confounding environmental factors. Therefore, the quantitative relationship between bacterial interactions and carbon source diversity remains largely unexplored.

To understand how carbon source diversity affects bacterial interaction classes, we analysed 896 interactions among 28 synthetic bacterial pairs, composed of eight species isolated from diverse habitats, under 32 different environments with varying carbon source diversity (**Fig. 1A, B**). Our results indicated that a broad range of interaction classes, from cooperative to competitive, were observed in single-carbon-source environments, whereas interactions shifted to competitive as carbon source diversity increased. Furthermore, detailed analyses suggested that these shifts were driven by intensified resource competition [34–37] in more complex media. This study offers new insights into the environmental determinants of bacterial interactions, especially the role of carbon source diversity. Notably, our findings suggest that bacterial interactions can be manipulated through environmental modifications focusing on carbon source diversity, providing strategies for managing bacterial communities in biotechnological applications [38, 39].

**Figure 1.**
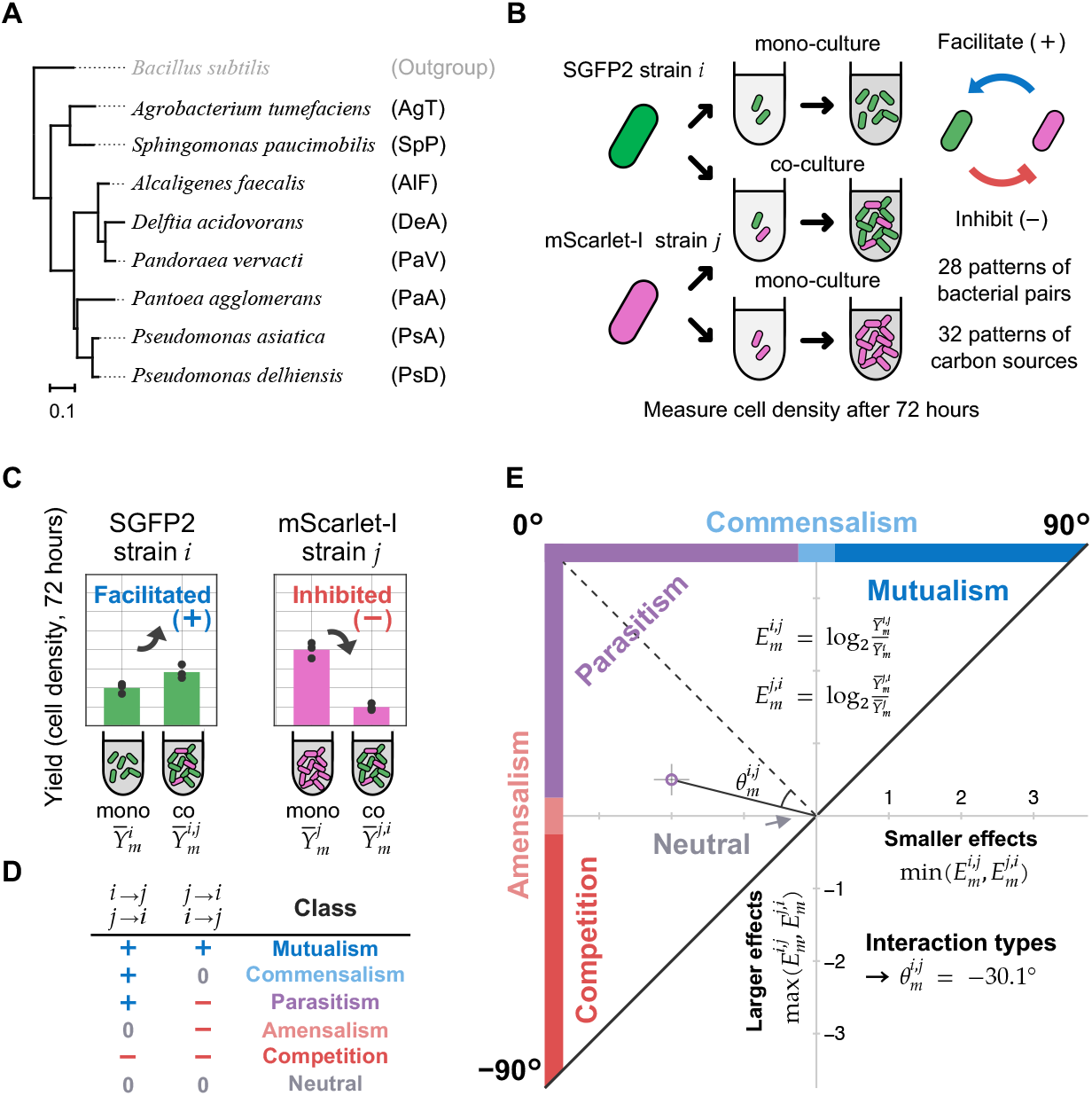
Overview of methods for measuring and analysing interspecies interactions. (**A**) Phylogenetic tree of the eight bacterial species used in this study, constructed based on 16S rRNA gene sequences. (**B**) Experimental setup for culturing experiments. Mono-cultures (single-species) and co-cultures (two-species) were performed for each bacterial pair (species *i* and *j*) in the growth environment *m*. Two unidirectional effects (from species *i* to *j* and from species *j* to *i*) in the co-culture were assessed by comparing fluorescent cell densities after 72 hours of incubation. (**C, D**) Classification of unidirectional effects (**C**) and bidirectional interactions (**D**). Fluorescent cell densities, which indicate growth yields (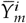 and 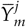 for mono-cultures; 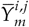 and 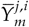 for co-culture), were used to assess unidirectional effects (facilitated [+], unaffected [0], or inhibited [–]) by comparing the growth (**C**). Bidirectional interactions were then categorised into six classes based on the combination of these unidirectional effects (**D**). (**E**) Quantification of interspecies interactions. The unidirectional effects were quantified as the log_2_ ratios of co-culture yields (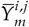 and 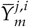) to mono-culture yields (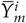 and 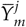), as described 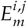 and 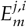. On a Cartesian plane, the smaller effects 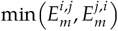 are plotted along the horizontal axis, while the larger effects 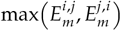 are plotted along the vertical axis. The interaction types 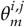 represent the angle that the line segment connecting this plotted point and the origin makes with the line where 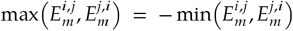. Error bars represent the standard deviation across biological triplicates.

## Materials and methods

### Culture media

All supplier information is provided in supplementary data (**Table S1**). All solutions were dissolved in pure water unless otherwise noted. To control the number of carbon source species in the growing environment, we used M9-based media prepared just before use. The media were prepared by dissolving the following components in 1.0 L of pure water: 100 mL of 10 × M9 salts solution (Na_2_HPO_4_ 67.78 mg/mL, KH_2_PO_4_ 30.00 mg/mL, NH_4_Cl 10.00 mg/mL, and NaCl 5.00 mg/mL, adjusted to pH 7.2 with 8.0 N NaOH), 1.0 mL of 1000 × Trace elements solutions (H_3_BO_3_ 24.8 µg/mL, MnCl_2_ 16.0 µg/mL, CoCl_2_ 7.2 µg/mL, (NH_4_)_6_Mo_7_O_24_ 3.4 µg/mL, ZnSO_4_ 2.8 µg/mL, and CuSO_4_ 2.4 µg/mL), 1.0 mL of MgSO_4_ solution (240.70 mg/mL), 1.0 mL of CaCl_2_ solution (11.10 mg/mL), and 1.0 mL of FeSO_4_ solution (1.52 mg/mL). Furthermore, carbon sources (**Supplementary methods S1, Table 1, Fig. S1**) and antibiotics were added as required (details are provided in each case). For agar plates, 15.0 g/L agar was also added. For construction of fluorescent-labelled strains, we used sterilised LB broth (25.0 g/L Difco™ LB Broth Miller), YENB broth (7.5 g/L Bacto™ Yeast Extract and 8.0 g/L Bacto™ Nutrient Broth), and NA broth (5.0 g/L Bacto™ Trypton and 3.0 g/L Bacto™ Yeast Extract) with antibiotics when needed (details are also provided in each case). Agar media also contained 15.0 g/L agar.

**Table 1.**
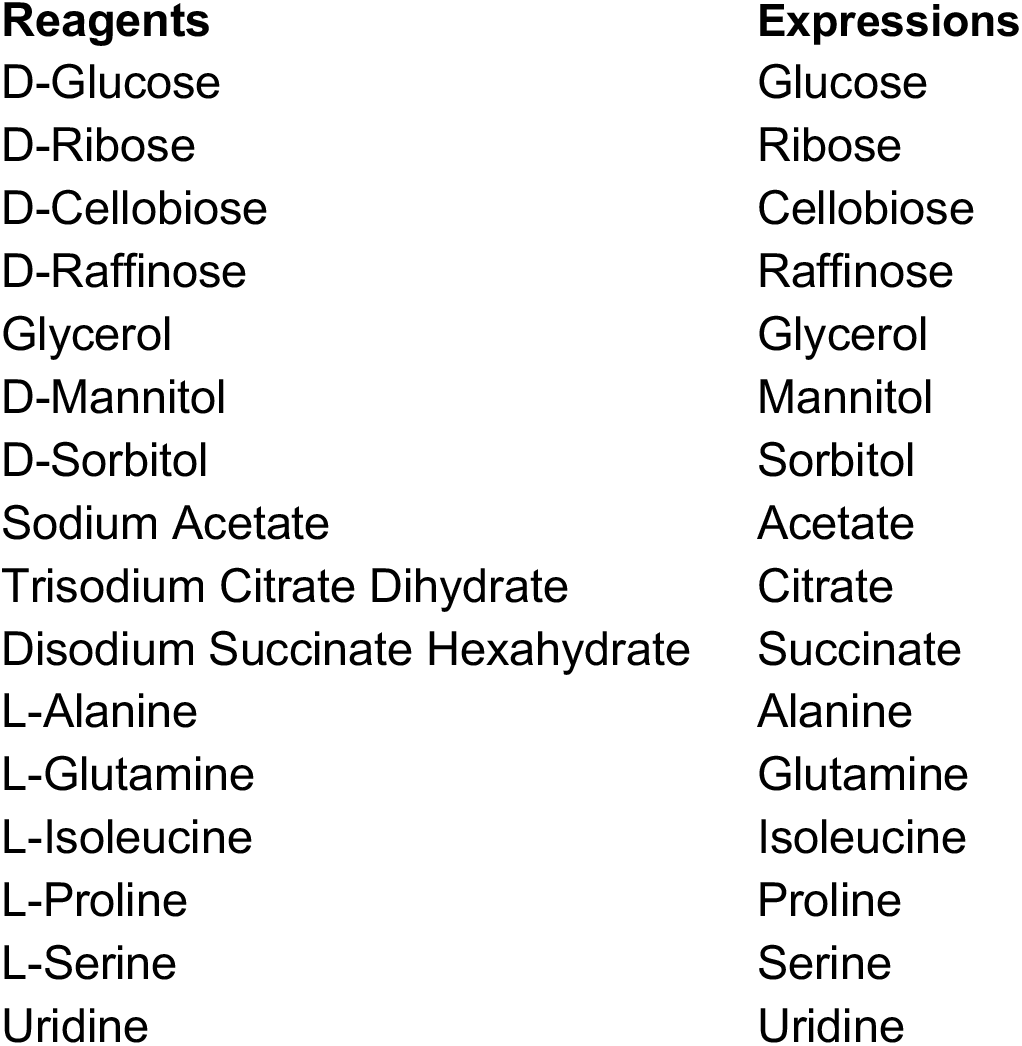
Carbon sources and their expressions.

### Bacterial strains

To select target species for our observations, we tested phenotypes of 192 bacterial species across five phyla, namely Pseudomonadota, Firmicutes, Actinobacteria, Bacteroidota, and Deinococcota (**Supplementary methods S2, Table S2**). Based on the selection, eight bacterial species were targeted for observation (**Figs. 1A** and **S1**). Notably, all selected species belonged to Pseudomonadota. To identify the species in two-species co-cultures, we constructed 14 fluorescent-labelled strains expressing green fluorescent protein (SGFP2) or red fluorescent protein (mScarlet-I) for each species (**Table S3**) [40]. These labelled strains grew distinctly in M9-based liquid medium, which contained 0.5 mg/mL glucose, 0.5 mg/mL succinic acid, and 20 µg/mL chloramphenicol. For abbreviations of bacterial species names, see **Fig. 1A**.

### Culturing assay

This step was performed using an automated culture system (Biomek NX span-8, Beckman Coulter, California, U.S.) in a clean booth, which was equipped with a plate reader (FilterMax F5, Molecular Devices, California, U.S.), plate shaker (StoreX STX44, LiCONiC, Mauren, Liechtenstein), and plate hotel (LiCotel LPX220, LiCONiC) [41]. First, 14 fluorescent-labelled strains were pre-cultured by inoculating frozen stocks into M9-based liquid medium adjusted to contain 0.5 mg/mL glucose, 0.5 mg/mL succinate, and 20 µg/mL chloramphenicol and incubating for 48 hours (200 µL, 96-well plate, 33°C, 500 rpm). After incubation, optical density at 595 nm (OD_595_) was measured, and appropriate dilutions were made according to the growth rate of each species based on these measurements. The diluted cultures were incubated again for an additional 18 hours under the same conditions. After the pre-culture, the strains were inoculated into M9-based liquid media, each containing 20 µg/mL chloramphenicol and a distinct carbon source composition from 32 different options. When inoculating, the fluorescent cell density of pre-cultures was measured by flow cytometry (see **Cell density measurement**). With the aim of investigating the growth limitations, we performed 14 strains mono-cultures in 16 distinct single-carbon-source environments (**Table 1**) at 0.5, 1.0, and 1.5 mg/ml. The results showed that for many species and carbon source combinations, the 1.0 mg/ml carbon sources were the limiting factors for growth (**Fig. S2**). For observing interactions, 14 strains one-species mono-cultures and 28 paired two-species co-cultures were performed in 32 different carbon source environments (**Figs. 1B** and **S3A**). These media included 16 single carbon sources, mixtures of 2, 4, 8, and 16 of carbon sources, and no carbon sources (**Fig. S3B**), with total carbon sources concentration adjusted to 1.0 mg/mL and equal contributions when mixed. Here, 20 µg/mL chloramphenicol was not assumed as the carbon sources for growth because it was sufficiently low compared to the other organic compounds. Pre-cultures were diluted to 2 000 cells/µL and 10 µL of them were inoculated into 190 µL of each mono-culture and 5 µL of them were added to 190 µL of each co-culture. We assumed that carbon sources in the pre-culture medium had negligible effects on interspecies interactions due to over 1 000-fold dilution. Cultures were shaken for 72 hours in which many combinations were saturated, with measuring optical density every three hours (200 µL, 96-well plate, 33°C, 500 rpm). After 72 hours, cultures were stored at −80°C for fluorescent cell density measurement.

### Cell density measurement

To measure fluorescent cell density, a portion of culture medium was mixed with an appropriate amount of standard fluorescent particle solution (Fluoresbrite® Polychromatic Red Microspheres 1.0 µm, Polysciences, Pennsylvania, U.S.) and PBS (KH_2_PO_4_ 144 µg/mL, NaCl 9 000 µg/mL, Na_2_HPO_4_ 421 µg/mL, pH 7.4), with the concentration of fluorescent particles known. Information on each particle in each diluted sample was collected by flow cytometry (FACS Aria™ III Cell Sorter, BD, New Jersey, U.S.; flow rate: 1.0, measurement time: one minute). Fluorescence information at 530 ± 15 nm wavelength emitted by the excitation light at 488 nm (green fluorescence in **Fig. S4**) and at 582 ± 7.5 nm wavelength emitted by the excitation light at 561 nm (red fluorescence in **Fig. S4**) were used for strains identification. The fluorescence information at 670 ± 7 nm wavelength emitted by the excitation light at 561 nm wavelength and the side scatter information were also used to identify the standard fluorescent particles. The cell density for each of the two fluorescent-labelled cell types was obtained by calculating the relative concentration to the standard fluorescent particles. The cell density of green or red fluorescent cells was set to 4.57 × 10^5^ cells/mL if it was lower than this threshold. This threshold was determined based on the 90th percentile of the total green and red fluorescent cell density observed in co-culture and mono-culture without carbon sources.

### Identification of the interaction classes

The interspecies interactions were classified for each combination of growth environment and bacterial pair. Following the method used in a previous study [31], we identified two unidirectional effects that occurred between the two species in the co-culture and classified the bidirectional interaction classes based on the combination of the two unidirectional effects. Specifically, we calculated the ratio of the fluorescent cell density of focal bacterial species in the co-culture at 72 hours to the average fluorescent cell density of the same species in mono-culture with the same carbon sources. This ratio is expressed as 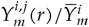, where 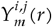 represents the fluorescent cell density for bacterial species *i* at 72 hours when the *r* -th replicate was co-cultured with bacterial species *j* in growth environment *m* (*r* ∈ {1, 2, 3}), and 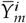 indicates the average cell density when focal species *i* was mono-cultured in the same environment *m*. Next, two-tailed t-tests were performed on the null hypothesis that the average of these values is equal to 1. Then, a multiple comparison using the Benjamini-Hochberg method [42] was conducted (False Discovery Rate [FDR]: 0.05). When the average of these ratios was significantly greater than 1, the unidirectional effect was considered positive (facilitated, +); less than 1, negative (inhibited, −); and when the null hypothesis was not rejected, it was considered to have no effects (unaffected, 0). Finally, the bidirectional interaction classes were categorised according to a widely recognised classification [5–7] based on the unidirectional effects (**Fig. 1D**).

### Quantification of the interaction types

Cooperation degree of interactions was quantitatively assessed as “interaction types”, based on the method used by Kehe et al. [33]. Interactions between bacterial species *i* and *j* in growth environment *m* were evaluated using equations (1) and (2):

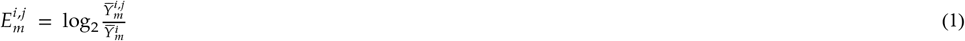

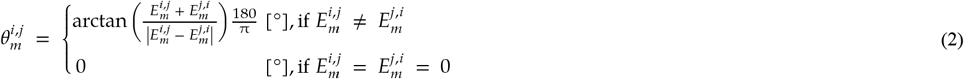

where 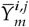 represents the average fluorescent cell density of species *i* at 72 hours of triplicate co-cultures with bacterial species *j* in growth environments *m* and 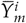 represents the average fluorescent cell density of species *i* at 72 hours of triplicate mono-cultures in growth environment *m*. 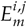 quantifies the unidirectional effects on species *i* when co-cultured with species *j* in the growth environment *m*. Similarly, 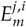 quantifies the unidirectional effects on species *j*. Then, we plot the smaller of these two values, 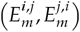, on the horizontal axis and the larger of these two values, 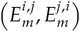, on the vertical axis in the Cartesian plane. In equation (2),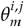 represents the angle 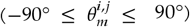 that the line segment connecting this plotted point and the origin makes with the line where 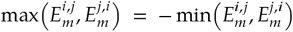 (dotted line in **Fig. 1E**), and was taken as a quantitative measure of the interaction types (higher values indicate more cooperative). When all values of 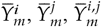 and 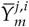 were at the lower limit of 4.57 × 10^5^ cells/mL, 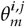 was set to 0°.

### Exploration of interaction determinants

To explore the determinants of the interspecies interactions, several statistical analyses were performed. Details are described in the supplementary information (**Supplementary methods S3–S7**). The raw data and analysis codes for R (version: 4.2.2) [43] is available on Zenodo [44] and GitHub (https://github.com/HirokiOnoGit/Carbon-sources-shape-bacterial-relations).

## Results

### Various interaction classes were observed in single-carbon-source environments

In this study, we analysed bacterial co-cultures in laboratory environments to explore the relationship between carbon source diversity and interspecies interactions. We used eight bacterial species isolated from various environments (**Table S2**), each labelled with either green (SGFP2) or red (mScarlet-I) fluorescent proteins [40], and paired them in combinations (**Figs. 1A, B**, and **S3A**). Cultures were grown in M9-based liquid media with 31 distinct carbon source compositions derived from 16 organic compounds, including sugars, alcohols, carboxylates, amino acids, and nucleic acids. These carbon sources were selected based on previous studies [33] and, due to operational constraints, were randomly combined, with up to 16 carbon sources per mixture (**Fig. S3B**). The total concentration of carbon sources was set at 1.0 mg/mL, which was found to be a level that restricted growth in many combinations of strains and single carbon sources (**Fig. S2**). When multiple carbon sources were included, they were mixed in equal amounts by mass. Additionally, cultures without any carbon source were included as controls. We conducted both mono-cultures and two-species co-cultures, measuring OD_595_ every three hours with a plate reader. After 72 hours, a timepoint sufficient for growth saturation in most combinations, fluorescent cell density was measured using flow cytometry. This allowed us to classify and quantify interspecies interactions statistically by comparing the growth of mono- and co-cultures under the same environmental conditions, as detailed in **Materials and methods** (**Fig. 1C–E**).

In single-carbon-source environments, various interaction classes were observed (**Fig. 2A, B**). The observed interaction classes were categorised as follows: cooperative interactions (mutualism and commensalism) accounted for 36.4%, exploitative interactions (parasitism) for 13.6%, competitive interactions (competition and amensalism) for 33.5%, and no interactions (neutral) for 16.5%. To reveal the factors that contributed to this wide variety of interspecies interactions, we then recapitulated observed interactions for each growth environment and bacterial pair as described below.

**Figure 2.**
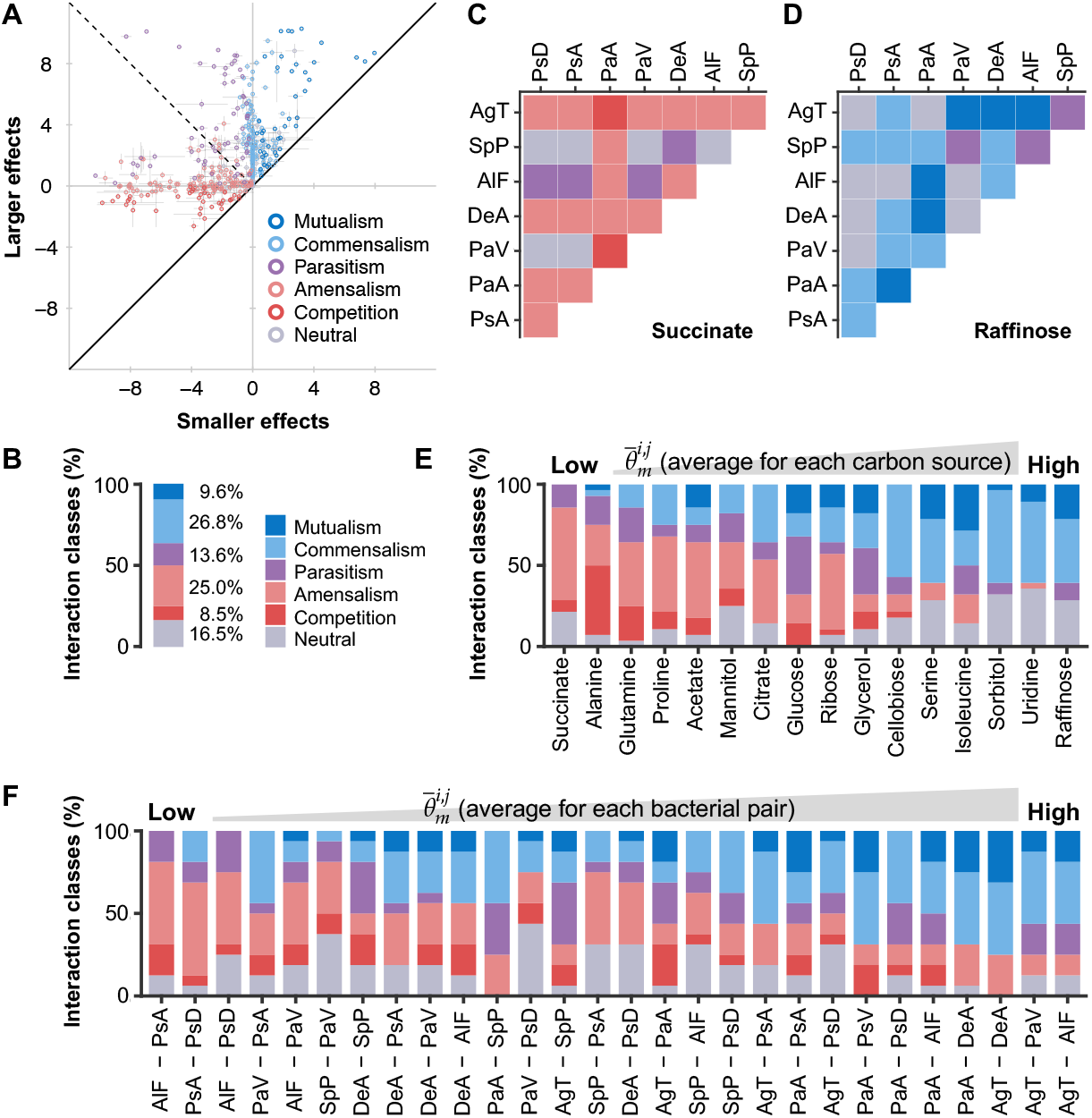
Diverse interspecies interactions in single-carbon-source environments. (**A**) Interspecies interactions for 28 bacterial pairs across 16 single-carbon-source environments. The smaller effects 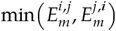 are plotted along the horizontal axis, while the larger effects 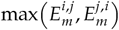 are plotted along the vertical axis. Colours represent different interaction classes. Error bars represent the standard deviation across biological triplicates. (**B**) Percentage bar chart of interaction classes observed in single-carbon-source environments, corresponding to data in panel **A**. (**C, D**) Classification of interspecies interactions when succinate (**C**) or raffinose (**D**) was provided as the single carbon source. Colours at the intersections of the vertical (species *i*) and horizontal (species *j*) axes represent the interaction classes between species *i* and *j*. (**E, F**) Interaction classes separated by carbon sources (**E**) or bacterial pairs (**F**). Bars in panel **E** are ordered by average interaction types 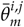 for each carbon source, while bars in panel **F** are ordered by average interaction types 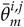 for each bacterial pair. See **Fig. 1A** for abbreviations of bacterial species names.

The interspecies interactions were significantly influenced by the provided carbon sources (chi-square tests [45], *p* = 1 × 10^−5^ [Monte Carlo estimated, 10^5^ iterations]; **Table S4**). To quantify interactions, we used the interaction types 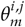 [33], representing the degree of cooperation for each bacterial pair comprising species *i* and *j* in the growth environment *m* (**Fig. 1E, Material and methods**). In brief, interaction types 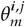 range from −90° to 90°, with higher values indicating more cooperative interactions. When 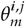 is −90°, both strains inhibit the growth of each other equally, whereas when 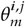 is 90°, both strains facilitate each other equally. Sorting the carbon sources by the average interaction types 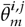 revealed that succinate promoted the most competitive interactions (**Fig. 2C**, leftmost column in **Fig. 2E**, 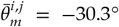), whereas raffinose led to the most cooperative interactions (**Fig. 2D**, rightmost column in **Fig. 2E**, 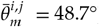). Previous research on genome-wide metabolic modelling has suggested that the biochemical category of carbon sources is one of the crucial factors explaining differences in bacterial interaction classes [23]. In contrast, Kehe et al. [33] pointed out that the biochemical category may not always explain interactions as well as previously expected. Given this, we performed a pairwise permutational multivariate analysis of variance (PERMANOVA) [46] on our data, following Kehe et al. and found no significant differences in interaction patterns within or between biochemical categories (FDR: 0.05; **Table S5, Fig. S5**). Additionally, hierarchical clustering of all interactions across bacterial pairs and carbon sources showed no clear grouping based on biochemical categories (**Fig. S6**). These findings suggest that the observed differences in interspecies interactions across different carbon sources are not driven by the biochemical category of carbon sources alone.

Interspecies interactions also varied depending on bacterial pairs. When comparing the average interaction types 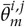 for each bacterial pair, *Alcaligenes faecalis* and *Pseudomonas asiatica* (AlF-PsA) formed the most competitive interactions 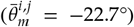, while *Agrobacterium tumefaciens* and *Alcaligenes faecalis* (AgT-AlF) formed the most cooperative interactions 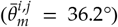 (**Fig. 2F**). Chi-square tests confirmed that bacterial pairs significantly influenced interaction classes (chi-square tests [45], *p* = 1.1 × 10^−4^ [Monte Carlo estimated, 10^5^ iterations]; **Table S6**). Nevertheless, many bacterial pairs displayed various interactions, ranging from competitive to cooperative, depending on the provided carbon sources (**Fig. 2F**). To determine whether certain species were predisposed to form cooperative or competitive interactions, we adapted the framework of gene set enrichment analysis (GSEA) [47]—originally developed for gene expression data—to our bacterial pair-interaction dataset (FDR: 0.05; **Table S7, Fig. S7**). No specific species were found to exhibit a significant bias toward either cooperative or competitive interactions, suggesting that interaction outcomes are affected by complex factors beyond species identity alone. Taken together, our findings indicate that both carbon sources and bacterial pairs influence interspecies interactions.

### Increasing the number of carbon source species led to more competitive interactions

To clarify how carbon source diversity affects bacterial interactions, we next examined the influence of increasing the number of carbon source species per medium. Carbon sources were mixed in equal mass amounts, set to a total concentration of 1.0 mg/mL, consistent with single-carbon-source environments. The supplementary data (**Table S8, Fig. S8**) presents all interspecies interactions across different carbon source and bacterial pair combinations. When categorising these results by number of carbon source species, we found that 33.5% of interactions were competitive in single-carbon-source environments (bottom row and leftmost column in **Fig. 3A**), whereas all interactions were competitive when the number of carbon source species reached 16 (bottom row and rightmost column in **Fig. 3A**). Furthermore, we confirmed that correlation between the number of carbon source species and the interaction types 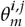 was negative (Spearman’s rank correlation: *ρ* = −0.41, *p* = 1.92 × 10^−33^; **Fig. 3B**). Based on these observations, we concluded that bacterial interactions tended to be more competitive as carbon source diversity increased.

**Figure 3.**
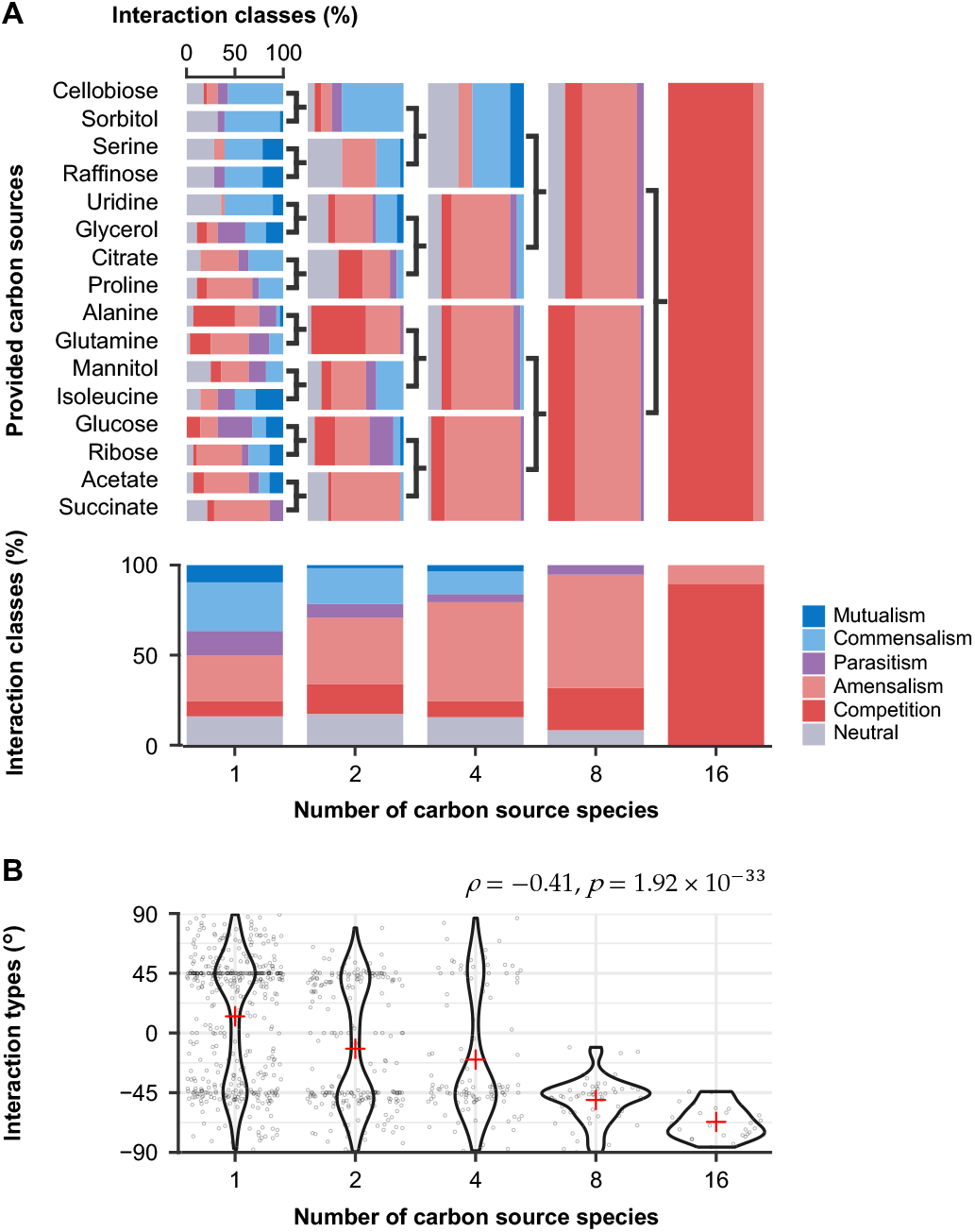
Relationship between carbon source diversity and bacterial interspecies interactions. (**A**) Percentage bar charts showing interaction classes in each environment. The upper panel displays horizontal bar charts categorised by provided carbon sources (rows) and the number of carbon source species (columns). Two-way symbols indicate that the carbon sources on the left were mixed and provided as a combination to the right. The lower panel presents vertical bar charts summarising interaction classes based on the number of carbon source species. The leftmost 16 horizontal bars are replots of **Fig. 2E**, while the leftmost vertical bar aligns with **Fig. 2B**. The datasets used for the rightmost horizontal and vertical bars are identical. (**B**) Violin plots with jittered points showing a negative correlation between the number of carbon source species and the interaction types 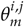. Cross-shaped points represent the average interaction types for each number of carbon source species.

### Competition is more frequent in environments where both species can grow easily

To understand how increased carbon source diversity led to more competitive interactions, we analysed the observations in detail. Potential mechanisms of competitive interactions include, for example, resource competition [34–37] and the production of growth inhibitors [48, 49]. In particular, in our set-up, the amount of provided carbon sources was found to be a growth constraint for most bacteria and carbon source combinations (**Fig. S2**). This leads us to speculate that the contest for carbon sources contributes as a major mechanism of competitive interactions. Based on this speculation, we proposed the hypothesis that increasing the number of carbon source species results in greater availability of carbon sources, which in turn leads to more frequent competitive interactions due to the intensified contest for available resources. We tested this hypothesis by evaluating two related sub-hypotheses: (1) increasing the number of carbon source species makes the environment easier for bacterial growth by improving the availability of carbon sources, and (2) competitive interactions occur more frequently in such “easy” environments for growth.

To test the sub-hypothesis (1), we first calculated 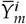, the average cell density of bacterial species *i* cultured triplicate in growth environment *m*. Notably, the distribution of 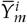 for each carbon source naturally formed two distinct clusters—one corresponding to cases with substantial growth and the other to cases with little to no growth. Given these data characteristics, we applied Ward’s clustering method [50] to formally distinguish these groups, designating them as “easy” environments where growth is possible and “hard” environments where growth is difficult (**Fig. S9**). Finally, we examined how the ease of growth varied with the number of carbon source species, and the results indicated that an increase in the number of carbon source species tended to make environments easier for bacterial growth (**Figs. 4A** and **S10**).

**Figure 4.**
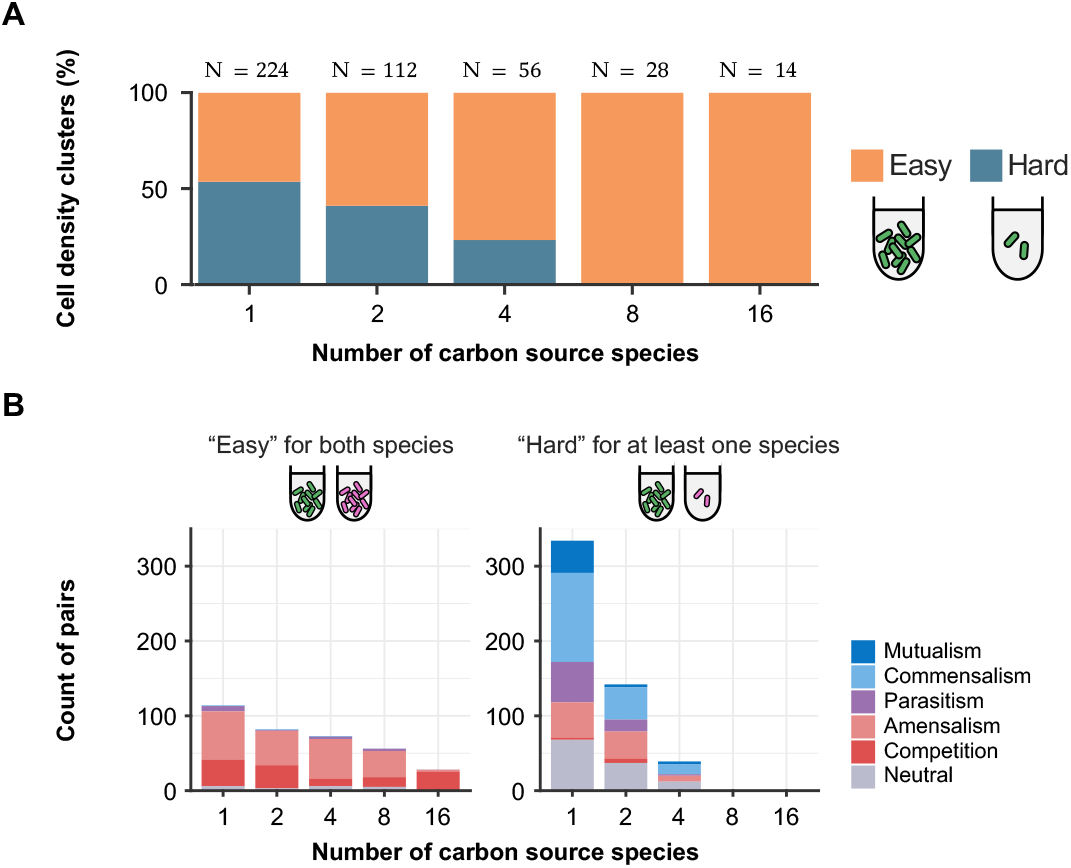
Competition is more frequent in environments that are easy to grow for both species. **(A)** Relationship between the number of carbon source species and growth ease in mono-cultures. Mono-cultures were categorised into “easy” or “hard” growth environments for each species based on growth yields, as detailed in the main text. Bars show the proportion of “easy” and “hard” mono-cultures for each number of carbon source species. Colours represent differences in growth ease. (**B)** Influence of growth ease on interspecies interactions. Bar charts show the distribution of interaction classes in environments where growth was “easy” for both species (left) and where growth was “hard” for at least one species (right). Colours represent different interaction classes.

Next, to evaluate the sub-hypothesis (2), we examined the interspecies interactions that occur in the “easy” environments where carbon sources are available for growth. We divided the bacterial interactions into two groups: those that occurred in an “easy” environment for both species, and those that occurred in a “hard” environment for at least one species of the pairs. The results showed that competitive interactions accounted for more than 82% of the interspecies interactions in the “easy” environment for both species (**Fig. 4B** left), regardless of the number of carbon source species (83.7%, 88.3%, 82.9%, 85.7%, and 100% when the carbon sources were 1, 2, 4, 8, and 16 species, respectively). In contrast, in environments that were “hard” for at least one species, a variety of interspecies interactions were formed, ranging from cooperative to competitive (**Fig. 4B** right). In order to statistically investigate whether specific interaction classes significantly accumulated in cases where the environments were either “easy” for both species or “hard” for at least one species, we performed an enrichment analysis. We analysed the data corresponding to environments with 1, 2, and 4 carbon source species, in which both “easy” and “hard” cases were identified, and found that competitive interactions were significantly accumulated in all of the cases where they were “easy” environments for both species (FDR: 0.05; **Table S9, S10**). Our analysis method did not detect competitive interactions when the average cell density in mono-culture is below 4.57 × 10^5^ cells/mL, which is the lower detection limit in our quantification. To rule out this methodological limitation, we performed the same analysis, excluding combinations where the average cell density in mono-culture is lower than this detection limit. The results confirmed that even after excluding these combinations, competition was still frequent in the “easy” environment (**Fig. S11**).

Based on the above, we conclude that both sub-hypotheses (1) and (2) are consistent with our observations. This suggests that the increased number of carbon source species leads to greater availability of carbon sources, which in turn drives the competitive interactions due to heightened contest for these resources.

### Growth in multi-carbon-source environments generally matches or exceeds the average growth when each carbon source is provided individually

Bacterial growth can be facilitated by different scenarios as carbon source diversity increases. In this study, we varied the number of carbon source species while keeping the total amount constant at concentrations expected to be fully consumed in most combinations. Under this condition, a straightforward expectation is that the growth yield of multi-carbon-source environments would be consistent with the average yield before mixing the corresponding single-carbon-source environments. Alternatively, some carbon source combinations may enhance growth by fulfilling nutrient requirements [51, 52], while others could suppress it due to carbon source toxicity [48, 49].

To reveal how increasing carbon source diversity influences bacterial growth, we compared growth yields in multi-carbon-source environments to those in the corresponding single-carbon-source environments. In two-carbon-source environments, we first calculated the average yield of bacterial species *i* cultured in a mixed environment with two carbon sources *m* and *n* across biological triplicates, denoted as 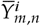. We then compared 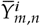 with the average yields of the same bacterial species in the corresponding single-carbon-source environments, 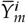 and 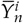. In 88% of cases, 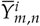 matched or exceeded the mean of 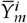 and 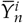 (two-tailed t-test, FDR: 0.05; **Figs. 5A** and **S12A**). Similarly, when 4, 8, and 16 carbon sources were provided, growth yields in multi-carbon-source environments generally matched or exceeded the average of the yields in the corresponding single-carbon-source environments (**Figs. 5B–D** and **S12B– D**).

**Figure 5.**
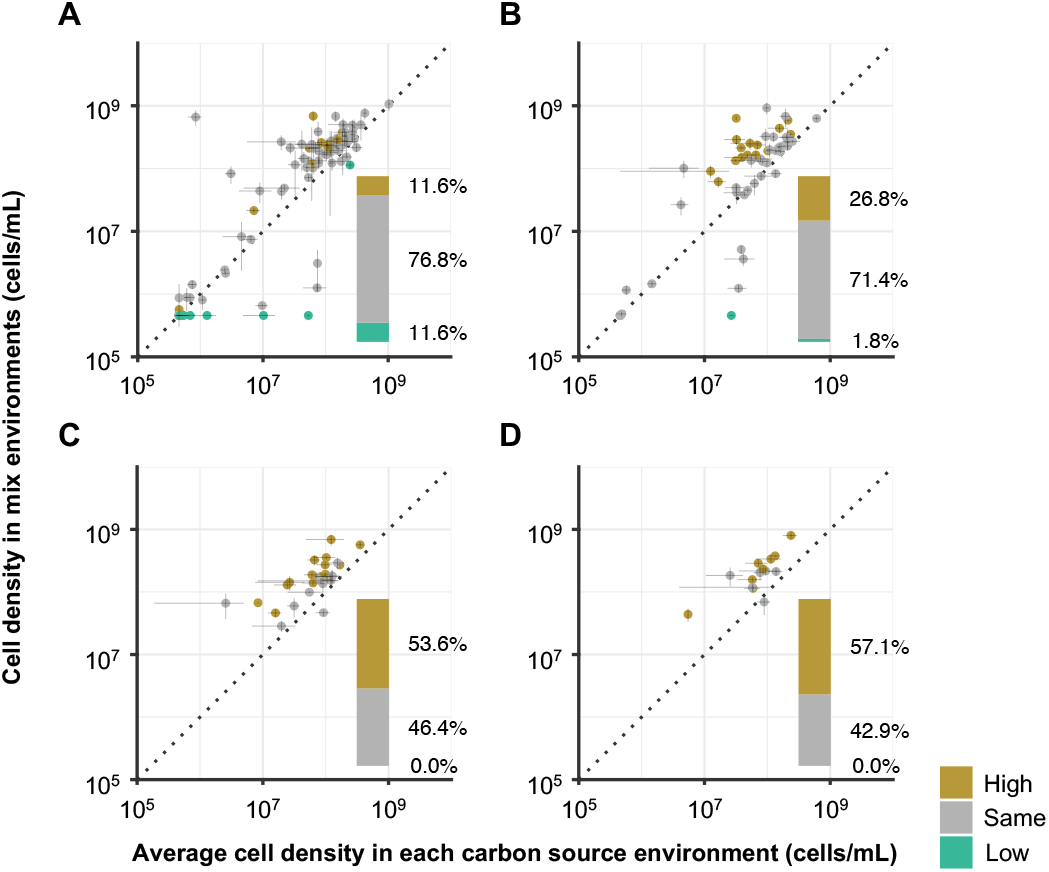
Relationship between growth yields in multi-carbon-source environments and the average yields in the corresponding single-carbon-source environments. The vertical axis represents the growth yields of species *i* in mono-cultures under multi-carbon-source environments, while the horizontal axis represents the average yields of the same species in mono-cultures when each carbon source was provided individually. The dashed lines represent the points where the values on the vertical and horizontal axes are equal. Colours denote three scenarios based on whether the vertical axis values are higher than, equal to, or lower than the horizontal axis values. Insets show the distribution of these three scenarios. Panels (**A**–**D**) correspond to environments containing 2, 4, 8, and 16 carbon source species, respectively.

While cell density serves as an important indicator of bacterial population growth, it does not fully capture biomass growth, including growth in cell size. To address this, we repeated the similar analysis using OD_595_ data as an alternative growth measure that accounts for growth in cell size [53]. The results were consistent, confirming that growth yield in multi-carbon-source environments was equal to or greater than the average yield in single-carbon-source environments (**Fig. S13**). As a more stringent test, we repeated the analysis after excluding environments where the carbon sources were likely not fully consumed. First, we identified these environments by comparing growth yields across different carbon source concentrations in single-carbon-source conditions (**Fig. S2**). Then, we excluded data from combinations where 1.0 mg/mL carbon sources were likely not fully consumed and repeated the analysis. The results derived from the refined dataset remained consistent with the previous results, indicating that incomplete carbon source consumption did not significantly influence the outcomes (**Fig. S14**). Overall, we conclude that bacterial growth in multi-carbon-source environments generally matches or exceeds the average growth in corresponding single-carbon-source environments. Thus, both simple additive effects and synergistic growth enhancement contribute to the facilitation of bacterial growth as carbon source diversity increases, likely leading to the intensified resource competition in co-cultures.

## Discussion

In this study, we observed bacterial interactions in laboratory environments and demonstrated that competitive interactions are more frequent in environments with higher carbon source diversity (**Fig. 3**). This finding underscores the critical role of not only the species combinations but also the growth environments in shaping interaction classes. Our findings also support the previous assertion by Kehe et al. [33] that competitive interactions are more likely in environments where bacteria can easily grow (**Fig. 4**). Additionally, we found that combining multiple carbon sources enhanced bacterial growth, either matching or exceeding the levels observed with individual carbon sources, indicating that greater carbon source diversity promotes bacterial growth (**Fig. 5**) [54, 55].

While our findings align with previous studies, we also observed some discrepancies. For example, Kehe et al. [33] reported that cooperative interactions accounted for 17% in single-carbon-source environments, whereas we found a significantly higher proportion of 36.4% (**Fig. 2B**). This discrepancy may be attributed to differences in the taxonomic diversity of bacterial species. We used a broader range of species spanning three classes, whereas Kehe et al. focused on species within a single class (**Fig. S15**) [33]. Since closely related species with similar metabolic networks are expected to engage in competitive interactions [34–37], the inclusion of distantly related species in our study likely resulted in fewer competitive interactions and a relative increase in cooperative interactions.

The relatively high taxonomic diversity in this study also provides new insights into how bacterial combinations influence interspecies interactions. Kehe et al. defined “metabolic distance” [33] based on bacterial growth data from 33 distinct carbon sources and found a positive correlation with the average interaction types (Spearman’s rank correlation: *ρ* = 0.75, *p* < 0.01). Similarly, Ona et al. [56] explored the relationship between phylogenetic distance and interspecies interactions, noting that competitive interactions were more common among closely related species, while cooperative interactions prevailed among distantly related species. However, our analysis including more cross-class pairs revealed a weaker correlation between metabolic or phylogenetic distance and interaction types compared to previous studies (**Figs. S16** and **S17**). For instance, the correlation between metabolic distance and interaction types was weaker (Spearman’s rank correlation: *ρ* = 0.07, *p* = 7.13 × 10^−3^), and no significant correlation was found when averaging the interaction types across bacterial pairs (Spearman’s rank correlation: *ρ* = 0.16, *p* = 0.41). These findings suggest that while metabolic or phylogenetic distances can explain interactions within a class, where resource competition dominates, they might be insufficient for predicting interactions across classes, where cooperation occurs more frequently [57].

Our study provides crucial insights into the determinants of bacterial interactions. Similar to our discussion on metabolic and phylogenetic distances, neither the information on the biochemical categories of carbon sources nor the presence of specific bacteria revealed clear patterns (**Tables S5** and **S7, Fig S5**). These results emphasise that the interaction types cannot be explained solely by these factors. On the other hand, it is remarkable that we discovered the carbon source diversity and ease of growth significantly influence bacterial interactions. Our findings suggest that factors influencing interactions in environments where bacterial growth is “easy”, such as competition for available carbon sources, could be one of the most critical determinants explaining the interaction types.

Although our experiments do not reveal the molecular mechanisms of the bacterial interactions, some previous studies are helpful in estimating them. For example, bacterial growth facilitation is known to be triggered by processes such as the exchange of metabolites [23, 58] and secretion of enzymes [59]. Regarding growth inhibition, it could stem from resource competition [34–37] and the secretion of inhibitory substances such as organic acids and antibiotics [48, 49]. We consider that the observed interactions reflect the net result of these processes occurring simultaneously. The relative contribution of these processes will vary according to environmental conditions and bacterial pairs. In order to reveal the details of specific molecular mechanisms, it would be useful to understand these processes in relation to the observed interactions, for example by omics analysis of transcripts and metabolites.

We also observed that increasing the number of carbon source species led to synergistic effects, where bacterial growth exceeded the average growth in individual carbon source environments (**Figs. 5** and **S12–S14**). These phenomena can be attributed to several mechanisms described below: the fulfilment of nutritional requirements [51, 52], the alleviation of specific metabolic stresses [60], and the optimisation of the metabolic network [61]. Each of these mechanisms may arise from increasing carbon source diversity and, in turn, enhance bacterial growth. Additionally, the effects due to mixing carbon sources has been actively studied as a phenomenon known as the priming effect [54, 55]. Although our study does not precisely identify the mechanisms, these previous studies may help elucidate the processes driving these synergistic effects.

To address whether the patterns observed in this and previous studies can be generalised to other environments or phylogenetic groups, additional investigations are required. In our experiments, we used relatively simple carbon sources, including compounds from the tricarboxylic acid cycle. This restriction may bias results in competition compared to macromolecules, which are expected to lead to cross-feeding [62, 63]. Similarly, we selected the bacteria observed in this study to satisfy the following two requirements: (1) the ability to grow in M9-based liquid medium supplemented with one of the following carbon sources—glucose, glycerol, succinate, or proline without visible aggregation; (2) the capacity for transformation (**Supplementary methods S2**). Our findings suggest that such culturable bacteria, which are expected to have access to a wider range of carbon sources [64–66], are more prone to competitive interactions. This indicates that competitive interactions are less common in communities containing unculturable bacteria. Natural environments, such as the gut [67] and soil [68, 69], are known to harbour more than thousands of bacterial species across multiple phyla, presenting greater ecological and phylogenetic complexity than the in vitro systems studied here. Further investigations are required to reveal whether the patterns we observed hold in such complex microbial communities.

Our findings offer novel strategies for developing techniques to regulate bacterial interspecies interactions. Despite the inherent complexity of bacterial growth systems, our study demonstrates that carbon source diversity facilitates bacterial growth in a surprisingly straightforward manner. Combined with the predominance of competitive interactions in environments where growth is “easy”, our observations suggest that information on growth in single-carbon-source environments could enable partial control of bacterial interactions. While progress has been made in regulating the strength of interspecies interactions [39, 70–72], controlling the interaction types remains largely unachieved. Our results indicate that modifying carbon source diversity can influence the ease of growth and, in turn, partially control the interaction types. This insight has potential applications, such as isolating unculturable bacteria [73, 74] or eliminating pathogenic bacteria [32, 75], by leveraging strategies that manipulate interspecies interactions. In addition, control over the interaction types may provide regulation over community function and species diversity [8–14]. Our study paves the way for the establishment of techniques to modulate interaction types, with the goal of developing practical approaches to control bacterial communities.

## Acknowledgments

We thank Dr. Tomoya Maeda (Hokkaido University and RIKEN, Japan) for providing the isolated environmental bacteria, and Dr. Atsushi Shibai (RIKEN) for advice on handling the automated culture system. The plasmids were gifts from Mitja Remus-Emsermann (Addgene plasmid #118486; http://n2t.net/addgene:118486; RRID: Addgene_118486) [40]. The *E. coli* S17-1 strain was purchased from National BioResource Project (NIG, Japan). The text was proofread using ChatGPT (GPT-4o).

## Author contributions

All authors contributed to the study design. H.O. conducted experiments, performed analyses, created visualisations, and wrote the original manuscript. All authors reviewed and edited the manuscript. S.T. and C.F. supervised the project and acquired funding.

## Data availability

The datasets generated during the current study are available in Zenodo [44]. The analysis codes are available on GitHub (https://github.com/HirokiOnoGit/Carbon-sources-shape-bacterial-relations).

## Supplementary data

supplementary-information - pdf file

supplementary-tables - xlsx file

