## supplementary-information for "Carbon source diversity shapes bacterial interspecies interactions"

### Title

### Short title

Carbon sources shape bacterial relations

### Authors

Hiroki Ono[1], Saburo Tsuru[2*], and Chikara Furusawa[2,3*]

### Affiliations

[1]Department of Biological Sciences, Graduate School of Science, The University of Tokyo, 7-3-1 Hongo, Bunkyo-ku, Tokyo 113-0033, Japan

[2]Universal Biology Institute, Graduate School of Science, The University of Tokyo, 7-3-1 Hongo, Bunkyo-ku, Tokyo 113-0033, Japan

[3]Center for Biosystems Dynamics Research, RIKEN, 6-7-1 Minatojima-minamimachi, Chuo-ku, Kobe 650-0047, Japan

### *Corresponding authors

Saburo Tsuru

Chikara Furusawa

**Contents**


**Supplementary methods S1** Reagent stocking.

**Supplementary methods S2** Selection and construction of bacterial strains.

**Supplementary methods S3** Statistical analyses of interspecies interactions.

**Supplementary methods S4** Analysis of mixing carbon sources impact.

**Supplementary methods S5** Phylogenetic distance calculation.

**Supplementary methods S6** Metabolic distance calculation.

**Supplementary methods S7** Data visualisation.

**Figure S1** Carbon source utilisation profiles of observed bacterial strains.

**Figure S2** The amount of carbon sources can limit growth.

**Figure S3** Bacterial pairs and carbon source composition.

**Figure S4** Identification of fluorescent-labelled cells.

**Figure S5** Interaction classes grouped according to the biochemical categories of carbon sources.

**Figure S6** A heat map showing all observed interactions in single-carbon-source environments.

**Figure S7** Species biases in interaction types.

**Figure S8** Diagrams showing the interaction classes observed in each combination of bacterial pairs and carbon sources.

**Figure S9** Determination of “easy” environments for each bacterial species.

**Figure S10** Average cell density for each combination of bacterial species and carbon sources.

**Figure S11** Interspecies interactions when combinations that do not grow during mono-culture are excluded.

**Figure S12** Growth in multi-carbon-source environments generally matches or exceeds the average growth when each carbon source is provided individually.

**Figure S13** Changes in optical density when carbon sources were mixed.

**Figure S14** Changes in cell density caused by mixing carbon sources in combinations where the provided carbon source is assumed to limit growth.

**Figure S15** Phylogenetic relationships with bacteria used in previous studies.

**Figure S16** Relationship between metabolic distance and interaction types.

**Figure S17** Relationship between phylogenetic distance and interaction types.

### Supplementary methods S1 Reagent stocking.

The added carbon sources and antibiotics were prepared by dissolving the sterilised −20°C stock solutions in pure water at the following concentrations: D-glucose: 25.00 mg/mL, D-ribose: 25.00 mg/mL, D-cellobiose: 25.00 mg/mL, D-raffinose hexahydrate: 29.46 mg/mL, glycerol: 25.00 mg/mL, D-mannitol: 25.00 mg/mL, D-sorbitol: 25.00 mg/mL, sodium acetate: 34.73 mg/mL, trisodium citrate dihydrate: 38.88 mg/mL, disodium succinate hexahydrate: 58.19 mg/mL, L-alanine: 25.00 mg/mL, L-glutamine: 25.00 mg/mL, L-isoleucine: 25.00 mg/mL, L-proline: 25.00 mg/mL, L-serine: 25.00 mg/mL, uridine: 25.00 mg/mL, chloramphenicol: 2.00 mg/mL, ampicillin sodium: 106.29 mg/mL, kanamycin sulfate: 60.12 mg/mL (Note: some solutes, such as chloramphenicol, have low solubility in pure water, so we ensured that there was no sedimentation occurring each time they were used).

### Supplementary methods S2 Selection and construction of bacterial strains.

**Step1: Growth phenotype selection**

Bacterial species that we study must be able to grow in liquid M9-based medium in mono-culture. Additionally, in order to measure cell density accurately by flow cytometry, we need them to avoid forming agglutination in the same medium. Furthermore, in order to distinguish between the two-species in co-culture, it is necessary to be able to construct strains that express fluorescent protein. First, phylogenetically diverse 192 bacterial species across five different phyla (Pseudomonadota, Firmicutes, Actinobacteria, Bacteroidota, and Deinococcota) were selected as candidates for this study (**Table S2**). These bacteria include environmental strains isolated by Dr. Tomoya Maeda from various environments and some laboratory strains. These bacteria were shaken and cultured in M9-based liquid medium with either glucose, glycerol, succinate, or proline added as a single carbon source at a final concentration of 1.0 mg/mL (200 µL, 96-well plate, 32°C, 800 rpm). After 96 hours of incubation, optical density at 595 nm ($OD_{595}$) was measured, and species that did not show a certain level of turbidity in any environment and those that formed visible aggregation in at least one environment were eliminated from the candidates.

**Step2: Transformation**

The 35 species that were not excluded by the above selection were subjected to transformation experiments to obtain fluorescent-labelled strains (**Table S2**, column "Transformation experiments": yes). The transformation was carried out by using four plasmids (pMRE132, pMRE-Tn5-132, pMRE135, and pMRE-Tn5-135), which have a wide host range, to each bacterial species. These plasmids were gifts from Mitja Remus-Emsermann (Addgene plasmid #118486; http://n2t.net/addgene:118486; RRID: Addgene_118486) [1]. The *Escherichia coli* strain S17-1 (phenotype in **Table S3**) transfers copies of the plasmids to other bacteria by conjugation [2]. In order to use this property, we first constructed four donor *E. coli* strains, each carrying one of the plasmids by electroporation. The *E. coli* S17-1 strain was purchased from National BioResource Project (NIG, Japan).

Next, we tried to transform 35 recipient bacteria by conjugation. The four donor *E. coli* S17-1 strains were grown in LB liquid medium containing 20 µg/mL chloramphenicol for 48 hours with shaking (2 mL, 5 mL microtubes, 37°C, 230 rpm). In addition, 35 recipient bacteria were cultured in LB liquid medium for 48 hours (200 µL, 96-well plate, 32°C, 800 rpm). Afterwards, to obtain cells in the late exponential growth phase, 200 µL of the culture medium of strain *E. coli* S17-1 was added to 20 mL of pre-warmed LB liquid medium containing 20 µg/mL chloramphenicol at 37°C and further cultured with shaking (20 mL, 50 mL in a centrifuge tube, 37°C, 115 rpm). Also, the 35 recipient bacteria were diluted in other LB liquid medium pre-warmed at 32°C and cultured again (200 µL, 96-well plate, 32°C, 800 rpm). Here, dilutions were made between 4-fold and 100-fold according to the time until the end of the exponential growth phase, which was measured in advance for each bacterial species. After six hours, when growth could be confirmed visually, *E. coli* S17-1 was centrifuged (25°C, 7 000 g, 2 minutes, 50 mL centrifuge tube). To remove antibiotics from the medium, the supernatant was discarded and the pellet was resuspended in 20 mL of LB medium without chloramphenicol; this washing step was repeated twice. After suspension in 20 mL of LB, 500 µL of the washed *E. coli* S17-1 sample was mixed with 500 µL of each of the 35 recipient bacterial cultures and the cells in the mixture were precipitated by centrifugation (25°C, 7 000 g, 2 minutes, 1.5 mL microtubes). The mixture was concentrated by removing the supernatant to a residual volume of 50 µL, and 10 µL each of the mixtures was spotted on NA agar medium without spreading and incubated at 32°C. After 24 hours, cells on NA agar medium were suspended in 100 µL of PBS ($KH_2PO_4$ 144 µg/mL, NaCl 9 000 µg/mL, $Na_2HPO_4$ 421 µg/mL, pH 7.4) and spread on M9-based agar medium containing 1 mg/mL of each carbon source (glucose, glycerol, succinate, and proline) and appropriate antibiotics, followed by incubation at 32°C for one to two weeks. The antibiotics and their concentrations were determined based on the phenotypes contributed by the plasmids and the previously examined drug resistance of each bacterial species (details are shown in **Table S11**). After confirming the expression of fluorescent protein, the cells were spread on a new agar medium of the same composition and repeated at least three times to obtain a single colony completely isolated from *E. coli* S17-1. The isolated fluorescent-labelled strains were cultured in M9-based liquid medium containing 1 mg/mL of each carbon source (glucose, glycerol, succinate, and proline) along with appropriate antibiotics. After 72 hours, 150 µL of the culture was added to 50 µL of 60% glycerol, and the mixture was frozen at −80°C for stocks.

Eight strains that successfully constructed fluorescent-labelled strains were selected for subsequent studies (**Table S2**, column “Interaction observation”: yes). Note that the species successfully transformed were limited to Pseudomonadota. Additionally, by comparing the growth in the same medium with that of the non-transformed strain of the same species under conditions that did not contain chloramphenicol, it was confirmed that the transformants could be selected without noticeable growth inhibition.

### Supplementary methods S3 Statistical analyses of interspecies interactions.

To find out whether the interaction classes varied significantly by single carbon source or bacterial pair, chi-square tests [3] were performed. These tests were performed by grouping the interaction classes by each carbon source or bacterial pair and examining whether the differences were by chance or not in all pairwise cases. Pairwise permutational multivariate analyses of variance (PERMANOVAs) [4] were performed using vegan package [5] to examine the relationship between the observed interactions and the biochemical categories of carbon sources. These analyses were carried out by dividing the carbon sources into sugars (monosaccharides, disaccharides, trisaccharides), sugar alcohols, carboxylate ions, amino acids, and nucleic acids to see if the interaction classes observed in each were significantly biased compared to the full data. We also performed applied gene set enrichment analysis (GSEA) [7] using clusterProfiler package [8], which is widely used for gene expression analysis, to identify biases in the interaction classes likely to form in each bacterial species. This analysis was based on examining the bias in the average rank of the interaction type for each bacterial pair. Furthermore, to investigate the relationship between the ease of bacterial growth and interspecies interactions, we performed clustering analysis for each bacterial species using Ward's method [9]. This analysis utilised average cell density obtained from mono-culture experiments across 32 environmental conditions, identifying bacterial species-environment combinations as “easy” or “hard” for growth. Based on the results of this classification, an enrichment analysis was also performed to see whether each interaction class had significant accumulation in each case. This analysis was performed on data from environments containing one, two, or four carbon sources, where we identified cases in which the environments were either easy for both species or hard for at least one species. The clusterProfiler package [8] was used in this analysis.

### Supplementary methods S4 Analysis of mixing carbon sources impact.

The effect of mixing carbon sources on bacterial growth was investigated using yield indicators, cell density or optical density. This effect was examined by comparing Group 1, which represents growth in multi-carbon-source environments, with Group 2, which represents the average of growth when each carbon source is provided individually, as follows:

$$\{Y^i_{1,2,\ldots,n}(r) \mid r = 1,2,3\} \qquad \text{Group1}$$

$$\left\{\frac{\sum_{k=1}^{n}\bar{Y}^i_k}{n} + \sqrt{\frac{\sum_{k=1}^{n}(u^i_k)^2}{n}}, \frac{\sum_{k=1}^{n}\bar{Y}^i_k}{n}, \frac{\sum_{k=1}^{n}\bar{Y}^i_k}{n} - \sqrt{\frac{\sum_{k=1}^{n}(u^i_k)^2}{n}}\right\} \qquad \text{Group2}$$

where $Y^i_{1,2,\ldots,n}(r)$ represents the yield in the $r$-th mono-culture of species $i$ with a mixture of carbon sources 1, 2, ..., $n$. Each $\bar{Y}^i_m$ represents the average yield in the triplicate mono-cultures of species $i$ with carbon source $m$, and $(u^i_k)^2$ represents the unbiased variance of the yield, reflecting the variability in the mono-culture experiments. To check for significant changes in the means for these two groups, two-tailed t-tests were performed and evaluated (FDR: 0.05).

### Supplementary methods S5 Phylogenetic distance calculation.

Phylogenetic trees were constructed to examine evolutionary relationships among bacterial species. PCR amplification of the 16S rRNA gene region was performed (primers 27F [5′-AGAGTTTGATCMTGGCTCAG] and 1492R [5′-CGGTTACCTTGTTACGACTT] were used), and each PCR product was sequenced by Sanger sequencing (Sanger Sequencing Service, Azenta, New Jersey, U.S.). Then, BLAST [10] searches were performed on the core nucleotide database on the web page provided by National Center for Biotechnology Information (NCBI) to find bacterial strains with homologous genes to the determined sequences, and the full-length sequence of 16S rRNA gene was obtained. In addition, *Bacillus subtilis* was included as an outgroup. The sequences were aligned by MUSCLE [11] using the default parameter settings of the sequence analysis software MEGA11 [12, 13]. Then, the phylogenetic tree was constructed by using maximum likelihood method and general time reversible model [14]. A discrete gamma distribution was used to model evolutionary rate differences among sites (5 categories, parameter: 0.3137). The rate variation model allowed for some sites to be evolutionarily invariable (32.54% sites). All positions containing gaps and missing data were eliminated. Additionally, full-length 16S rRNA sequences of bacterial species from Kehe et al. [15] were similarly obtained and analysed for comparison. The sequence details are provided in supplementary data (**Table S12**).

### Supplementary methods S6 Metabolic distance calculation.

Metabolic distance [15] was calculated to quantify metabolic similarity between bacterial strains based on their growth across different carbon source environments. First, the average cell density for each strain was obtained from triplicate mono-cultures under 32 different carbon source conditions. The common logarithm of these values was taken, and a background value of $\log_{10}(4.57 \times 10^5)$ was subtracted for baseline correction. Next, for each strain, the resulting values were normalised by dividing by the maximum value observed for that strain. Finally, the Euclidean distance between strains was calculated using these normalised values across the 32 carbon source environments, providing a quantitative measure of metabolic distance.

### Supplementary methods S7 Data visualisation.

For data visualisation, ggplot2 package [16] and ggpubr package [17] were used to run the R codes.

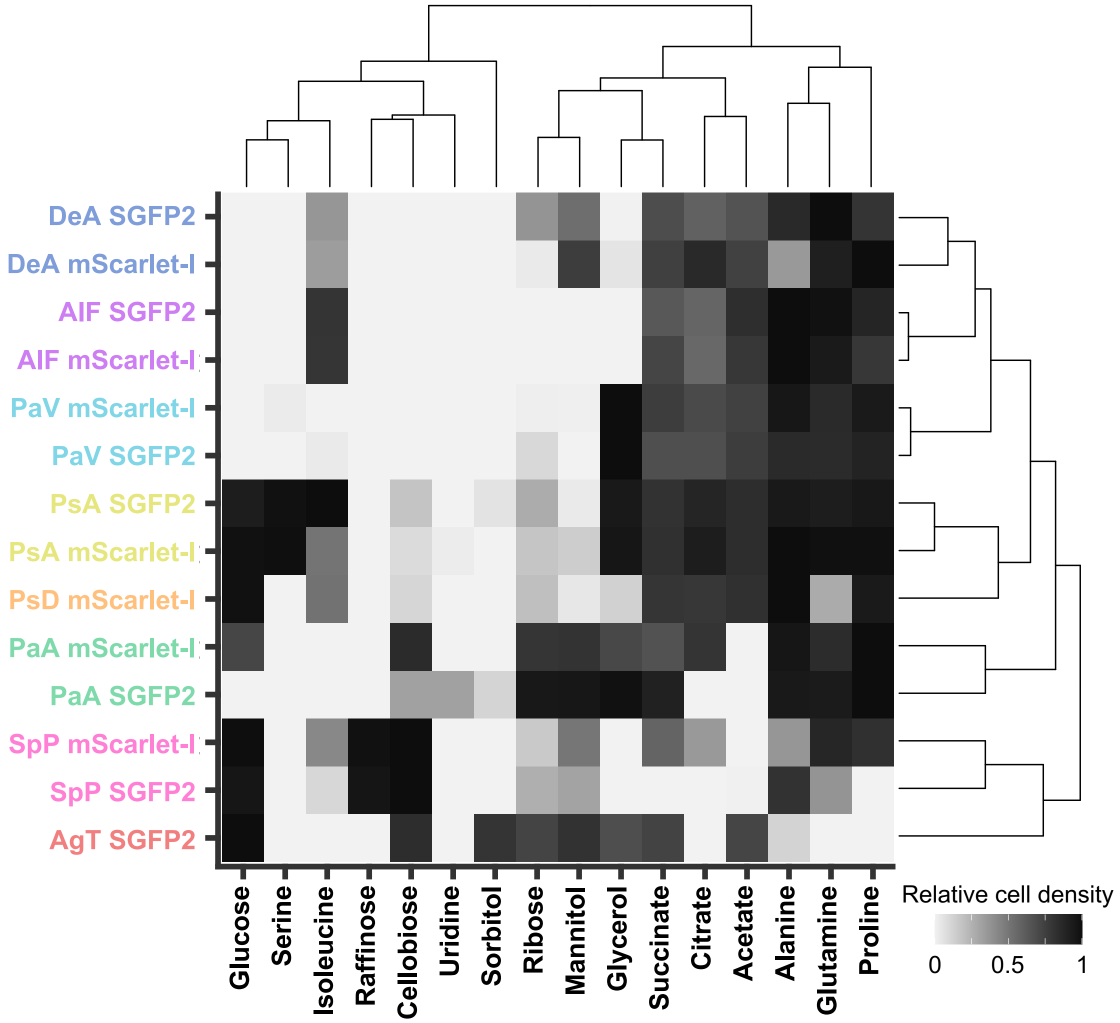


**Figure S1** Results of hierarchical clustering for the carbon source utilisation profiles. Gray scale of each grid shows the relative value calculated as the ratio of the cell density for each strain to the highest cell density observed among the carbon sources. Here, the relative values were calculated using the common logarithm of cell density minus background. The dendrograms represent the Euclidean distances, with bacterial strains and carbon sources arranged accordingly.

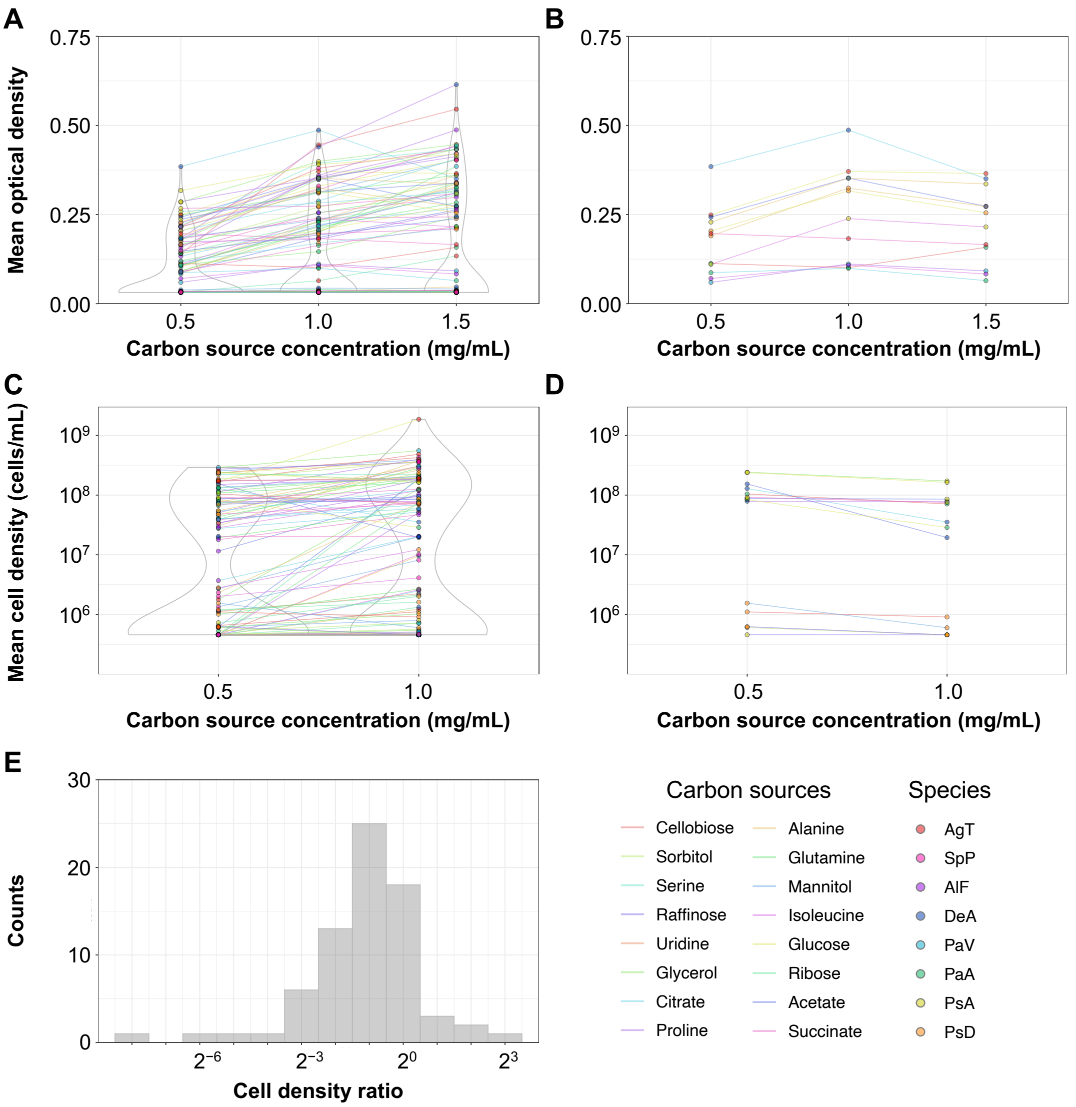


**Figure S2** Relationship between the concentrations of carbon sources and growth yields. (**A**, **B**) Optical density after 72 hours of mono-cultures for eight species in single-carbon-source environments with three different concentrations of carbon sources (0.5, 1.0, and 1.5 mg/mL). The horizontal axis represents the carbon source concentration in the growth environment, while the vertical axis represents the average optical density across triplicate mono-cultures for bacterial species $i$ with the carbon source $m$. Data points for the same species and carbon source combinations are connected by a straight line across different concentrations of carbon sources. **B** represents combinations where the optical density exceeded 0.05 at any carbon source concentration, and where a decrease in the average value of the optical density was observed despite increasing the carbon source in at least one of the concentration combinations (12 out of 128). (**C**, **D**) Cell density after 72 hours of mono-cultures for eight species in single-carbon-source environments with two different concentrations of carbon sources (0.5 and 1.0 mg/mL). The horizontal axis represents the carbon source concentration, and the vertical axis represents the average cell density across triplicate mono-cultures for bacterial species $i$ with the carbon source $m$. As above, data points for the same species and carbon source combinations are connected. **D** is a selection of combinations where the cell density exceeded $4.57 \times 10^5$ cells/mL at both carbon source concentration, and where the average value of cell density was found to decrease among combinations (14 out of 128). (**E**) A histogram showing the ratio of average cell density with 0.5 mg/mL of carbon source to the average cell density with 1.0 mg/mL, for each bacterial species and carbon source combination. Only combinations with cell density exceeding $4.57 \times 10^5$ cells/mL for any carbon source concentration are shown.

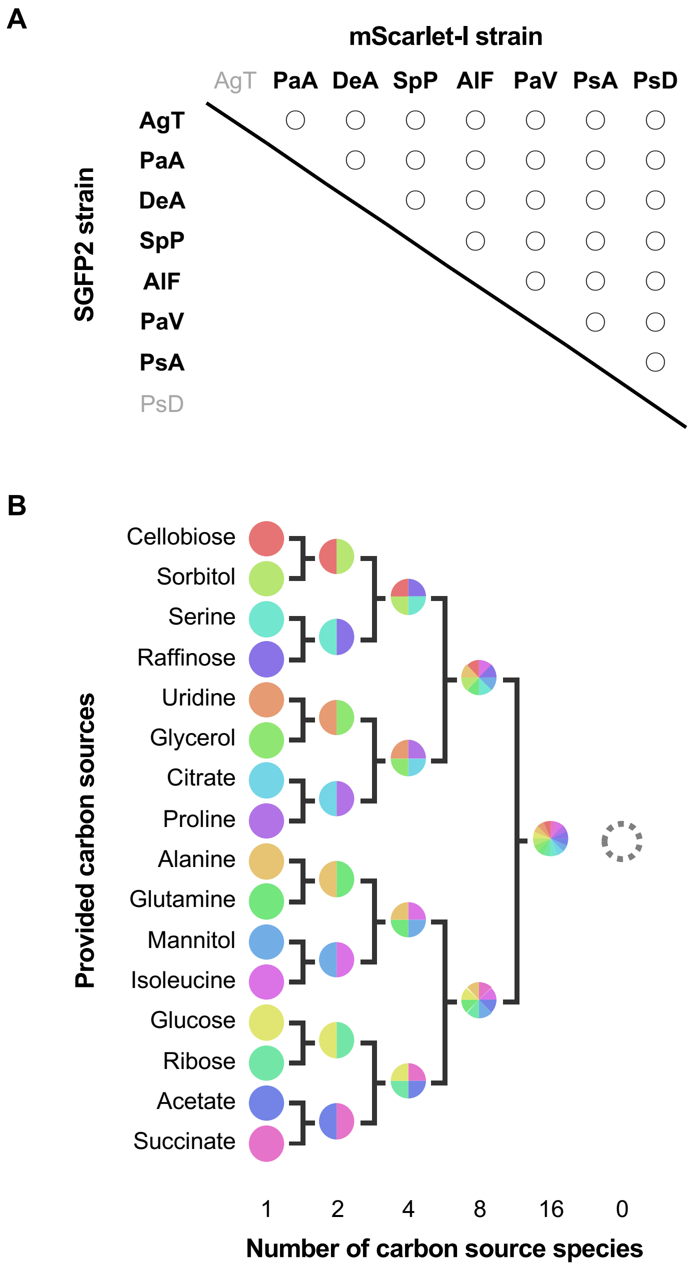


**Figure S3** Bacterial pairs and carbon source composition. (**A**) Strain combination of two-species co-culture. The 28 circles (○) represent pairs of co-cultures. The SGFP2 labelled strains are listed to the left and the mScarlet-I labelled strains above. The 14 bacterial strains represented in bold text were also mono-cultured to compare their yields with those of the co-cultures (SGFP2 labelled PsD and mScarlet-I labelled AgT were not mono-cultured). (**B**) Carbon source composition. The circles in all 32 ways represent the composition of the carbon source added to the M9-based liquid medium. Two-way symbols represent mixing of carbon sources. The combination of carbon sources was determined by following a tournament generated in a random order.

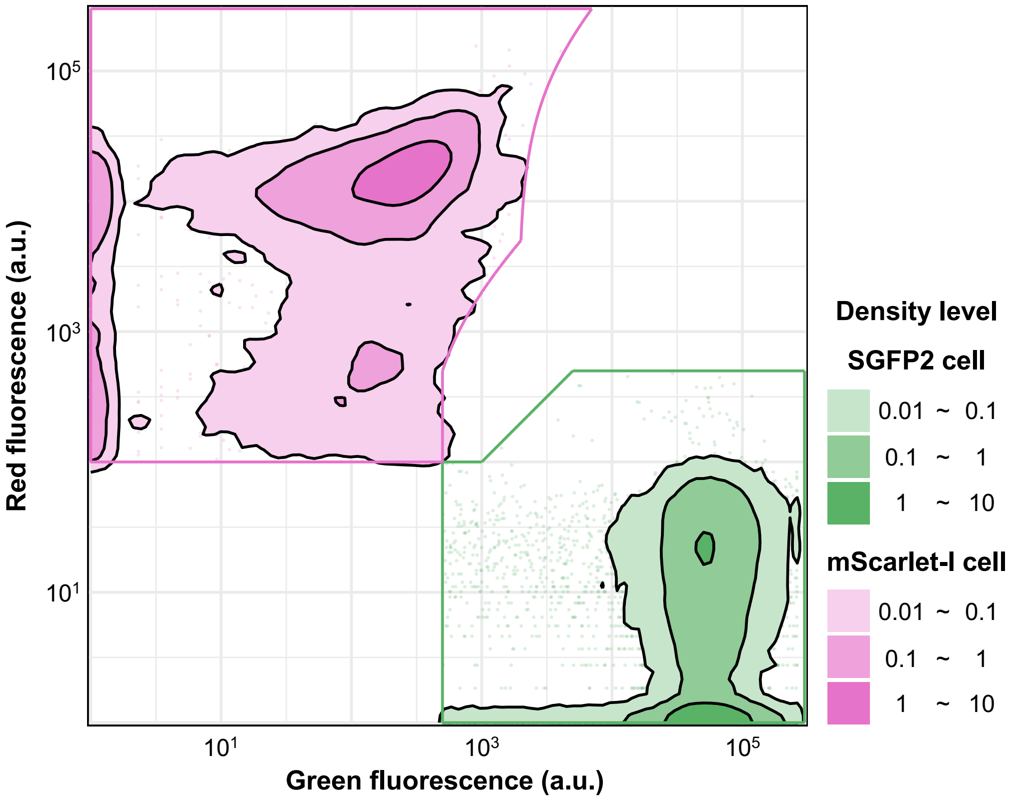


**Figure S4** Identification of fluorescent-labelled cells. The fluorescence information at a wavelength of 530 ± 15 nm emitted by an excitation light at 488 nm (green fluorescence) and at a wavelength of 582 ± 7.5 nm emitted by an excitation light at 561 nm (red fluorescence) is used to distinguish between SGFP2 labelled and mScarlet-I labelled cells. As a typical example, the analysis of a sample of SGFP2 labelled AgT and mScarlet-I labelled SpP co-cultured in a two-carbon-source environment, glucose and ribose, is shown. Only events discriminated as SGFP2 labelled or mScarlet-I labelled cells are shown here.

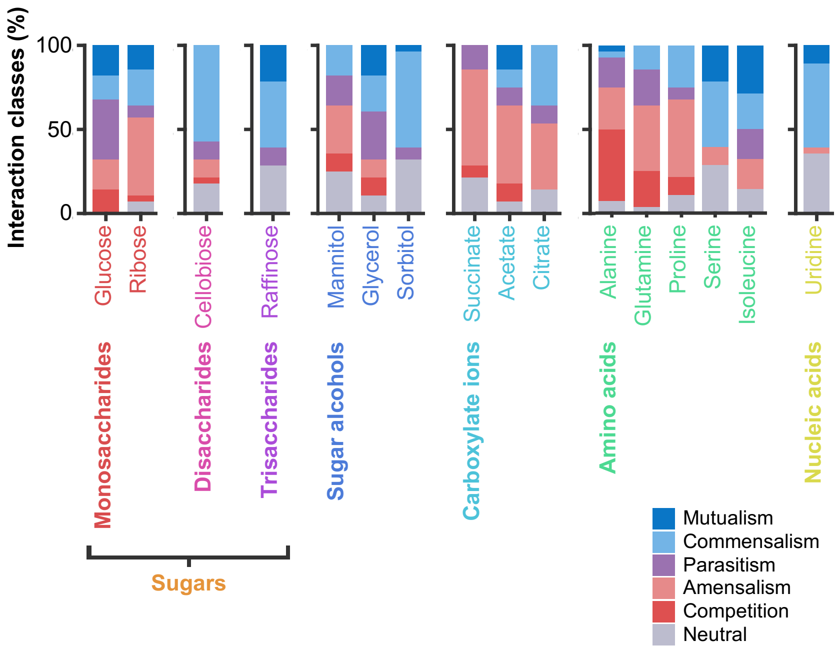


**Figure S5** Interaction classes grouped according to the biochemical categories of carbon sources. The interaction observed in single-carbon-source environments are shown. The colours of bar charts represent different interaction classes. The colours of labels indicate the biochemical categories of the carbon sources. This figure is a rearranged plot of **Fig. 2E**.

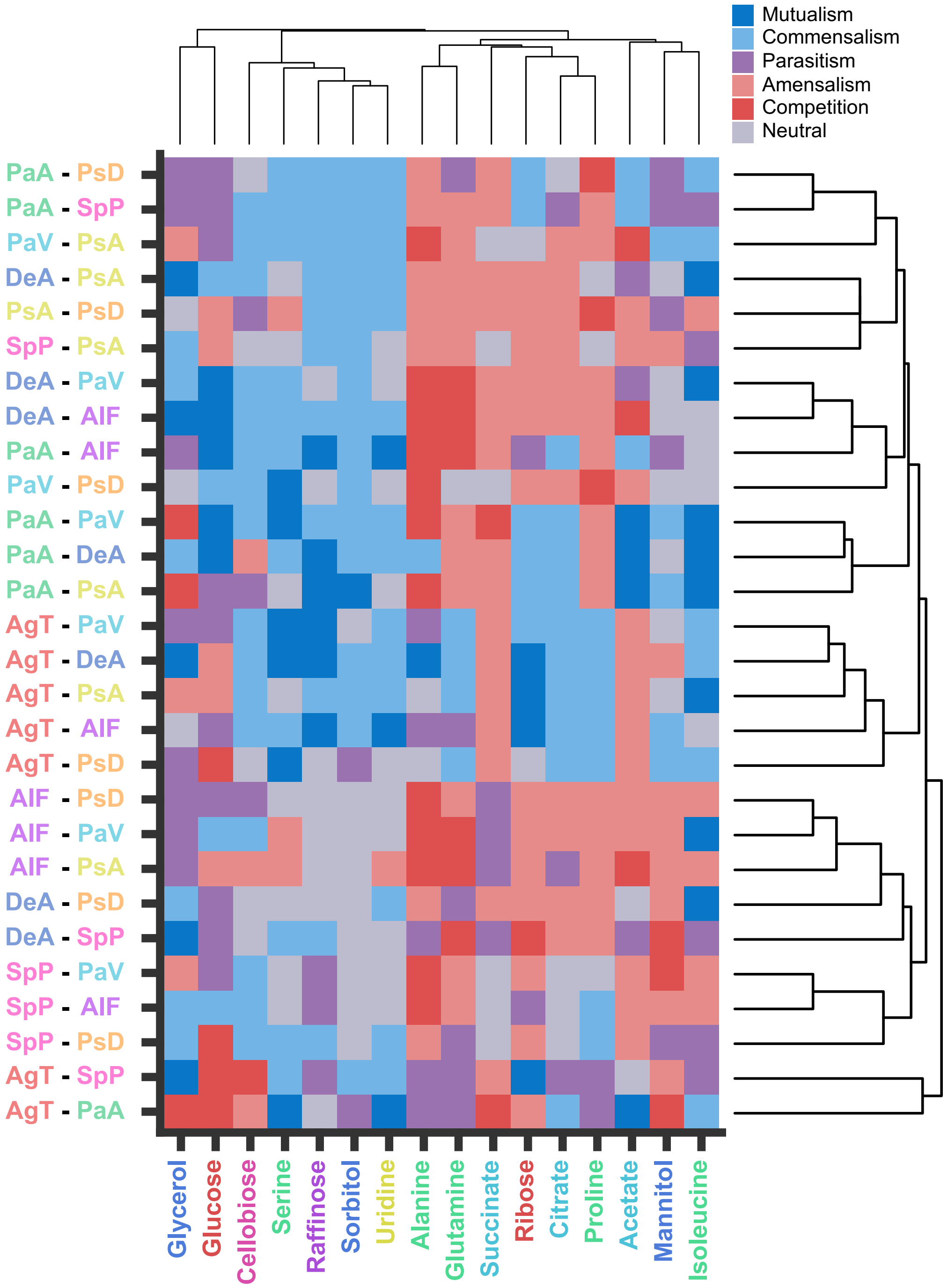


**Figure S6** A heat map showing all observed interactions in single-carbon-source environments. The colour of each grid represents the observed interaction class. The order of bacterial pairs and carbon sources is based on a dendrogram created from dice distances. The colour of each carbon source label refers to the biochemical categories. Each bacterium is coloured according to species.

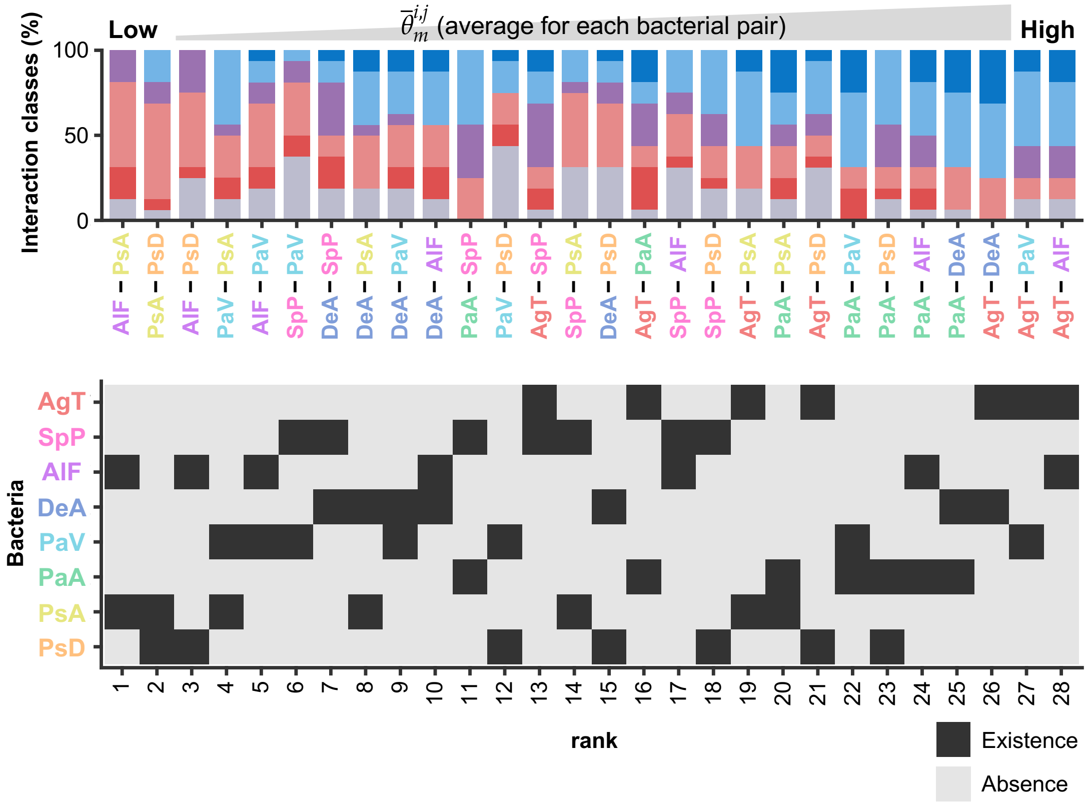


**Figure S7** Species biases in interaction types. The top panel is a rearranged plot of **Fig. 2F**, in which bacterial pairs are ranked by the average interaction types observed in single-carbon-source environments. The lower panel displays a heatmap indicating the presence or absence of each bacterial species in the pairs.

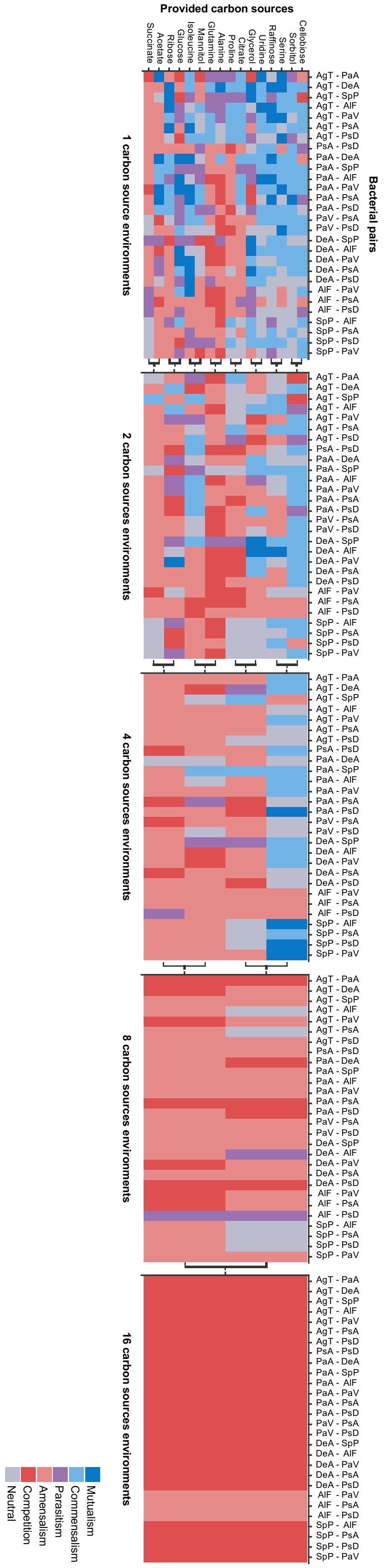

Provided carbon sources
Succinate
Acetate
Ribose
Glucose
Isoleucine
Mannitol
Glutamine
Alanine
Proline
Citrate
Glycerol
Uridine
Raffinose
Serine
Sorbitol
Cellobiose
Bacterial pairs
AgT - PaA
AgT - DeA
AgT - SpP
AgT - AIF
AgT - PaV
AgT - PsA
AgT - PsD
PsA - PsD
PaA - DeA
PaA - SpP
PaA - AIF
PaA - PaV
PaA - PsA
PaA - PsD
PaV - PsA
PaV - PsD
DeA - SpP
DeA - AIF
DeA - PaV
DeA - PsA
DeA - PsD
AIF - PaV
AIF - PsA
AIF - PsD
SpP - AIF
SpP - PsA
SpP - PsD
SpP - PaV
1 carbon source environments
2 carbon sources environments
4 carbon sources environments
8 carbon sources environments
16 carbon sources environments
Mutualism
Commensalism
Parasitism
Amensalism
Competition
Neutral

**Figure S8** Heatmaps showing the interaction classes observed in each combination of bacterial pairs and carbon sources. Colours represent different interaction classes. Two-way symbols represent mixtures of carbon sources. The figure for the case where one carbon source species was provided is a rearranged plot of **Fig. S6**.

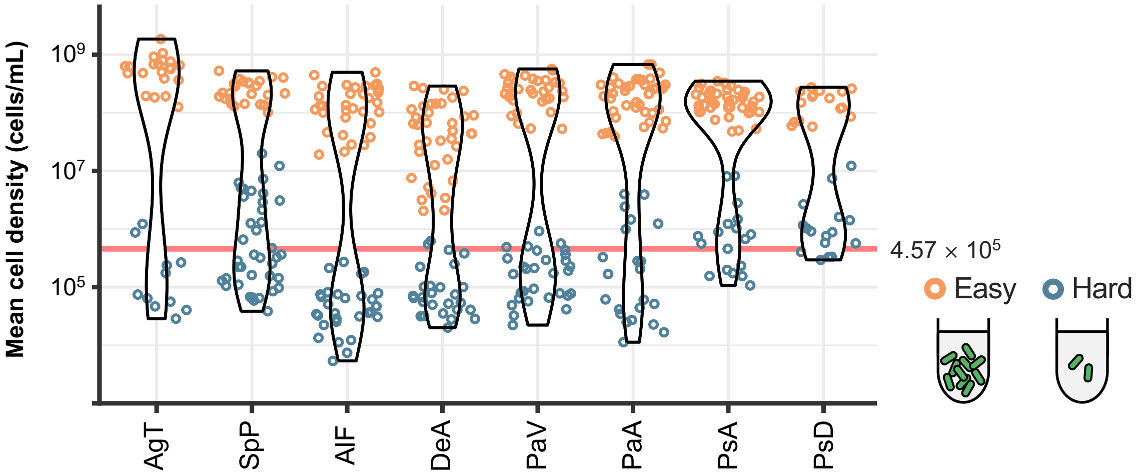


**Figure S9** Determination of “easy” environments for each bacterial species. Violin plots show the average cell density measured in 32 different growth environments for each bacterial species The colour of each point indicates clustering results, which are divided into two clusters, “easy” and “hard”.

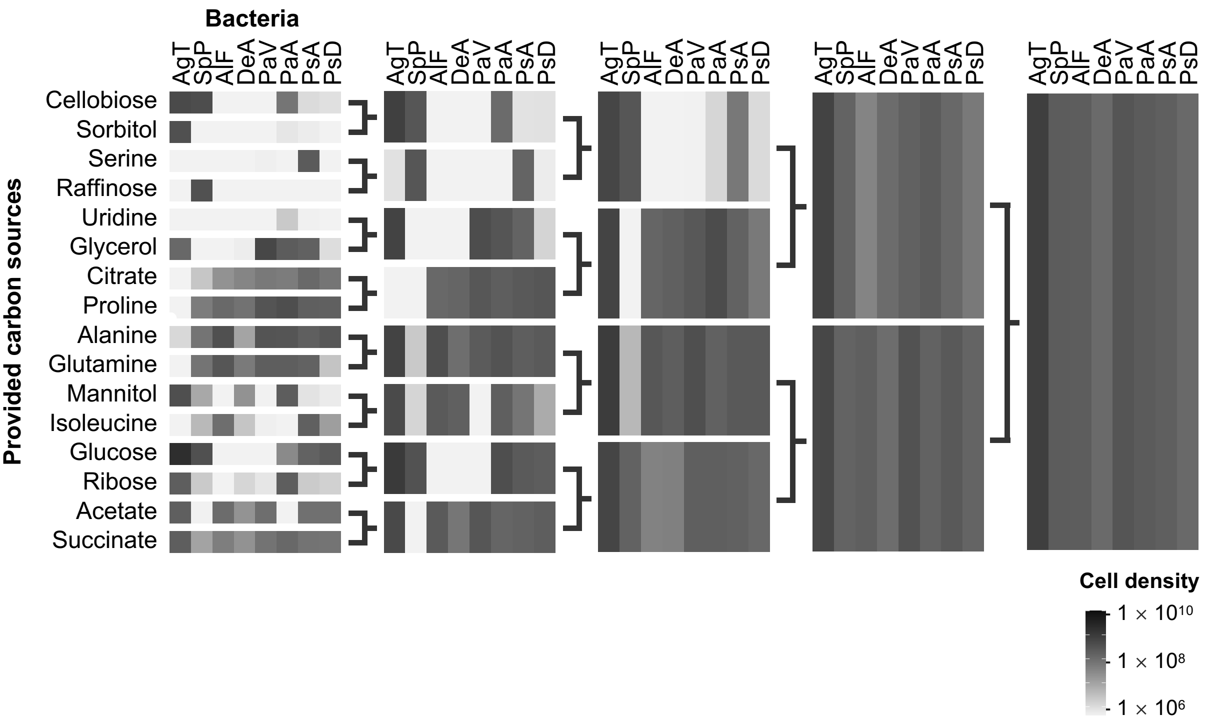


**Figure S10** Average cell density for each combination of bacterial species and carbon sources. For each bacterial species, the average cell density measured for each carbon source is indicated in greyscale. Two-way symbols indicate mixtures of carbon sources.

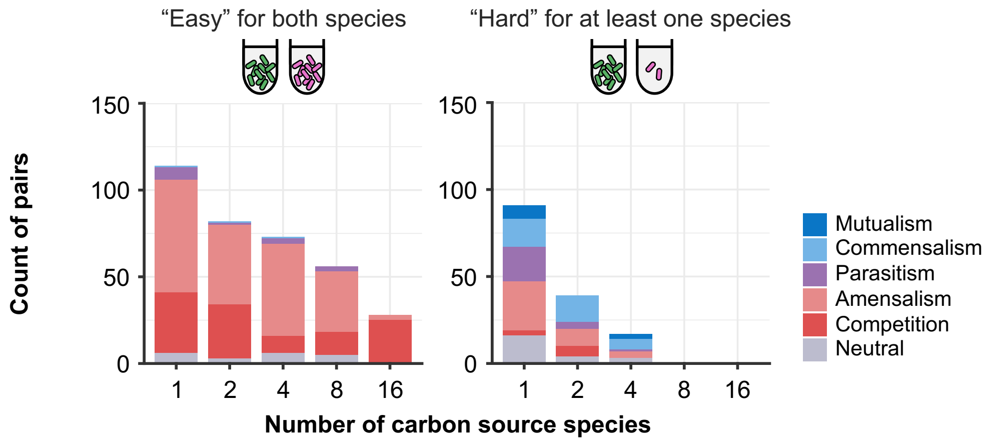


**Figure S11** Interspecies interactions when combinations that do not grow during mono-culture are excluded. Two histograms show the interaction classes when excluding combinations where the cell density obtained during mono-culture was below $4.57 \times 10^5$, the lower limit of measurement. As shown in **Fig. 4B**, the interactions were divided into those observed in environments that were “easy” for both bacterial species growth on the left and those observed in environments that were “hard” for at least one species on the right. The colours of bar charts represent different interaction classes.

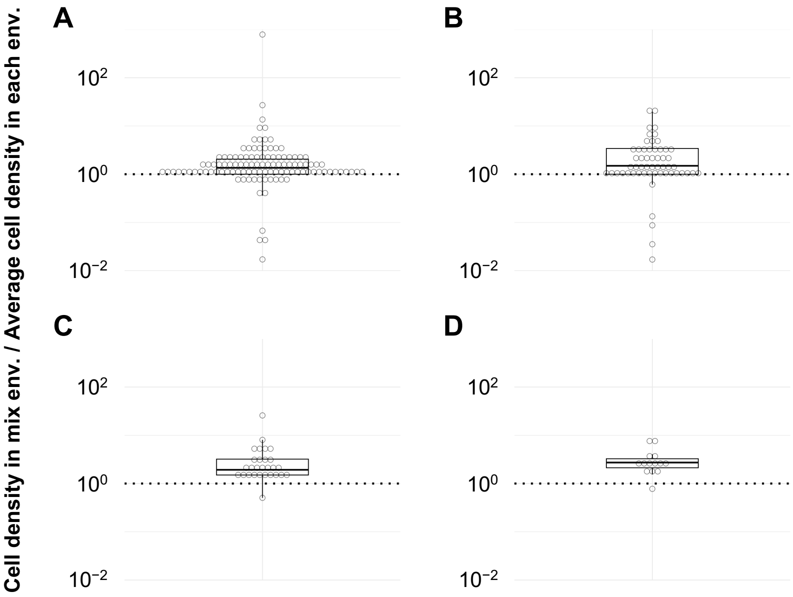


**Figure S12** Growth in multi-carbon-source environments generally matches or exceeds the average growth when each carbon source is provided individually. (**A**–**D**) Boxplots showing the ratio of the average cell density actually measured in multi-carbon-source environments to the average cell density when each carbon source was provided individually. Dotted lines indicate the case where the ratio is equal to $10^0$. **A**–**D** represent the cases where the number of carbon source species after mixing was 2, 4, 8, and 16, respectively.

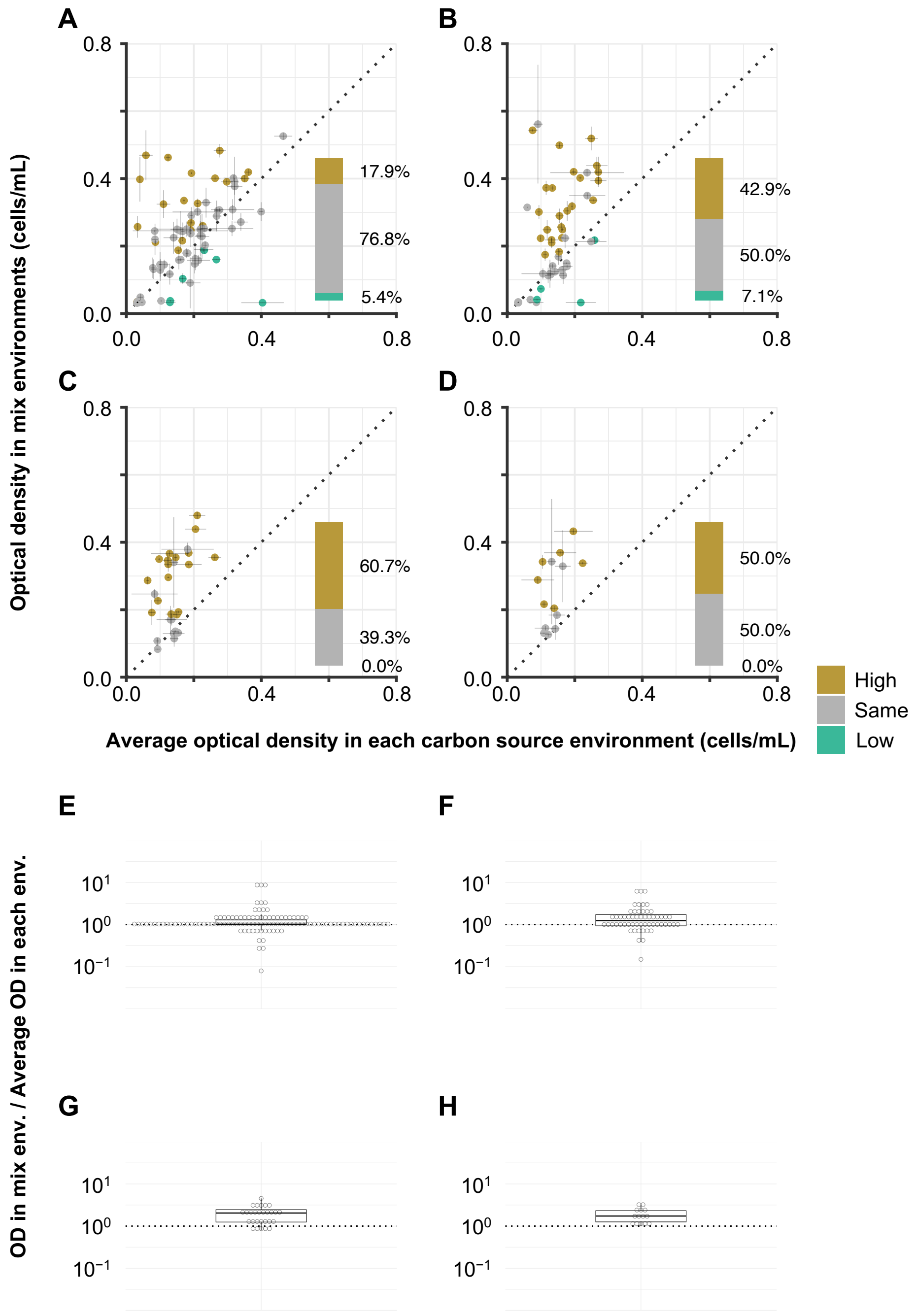


**Figure S13** Changes in optical density when carbon sources were mixed. (**A**–**D**) Similar to **Fig. 5**, scatter plots show the comparison of optical density of bacterial mono-cultures in multi-carbon-source environments with the average optical density when each carbon source was provided individually, coloured in the same way. **A**–**D** represent the cases where the number of carbon source species after mixing was 2, 4, 8, and 16, respectively. (**E**–**H**) Similar to **Fig. S12**, the ratios of the average optical density actually measured in the multi-carbon-source environments to the average optical density when each carbon source was provided individually are shown in boxplots. **E**–**H** also represent the cases where the number of carbon source species after mixing was 2, 4, 8, and 16, respectively.

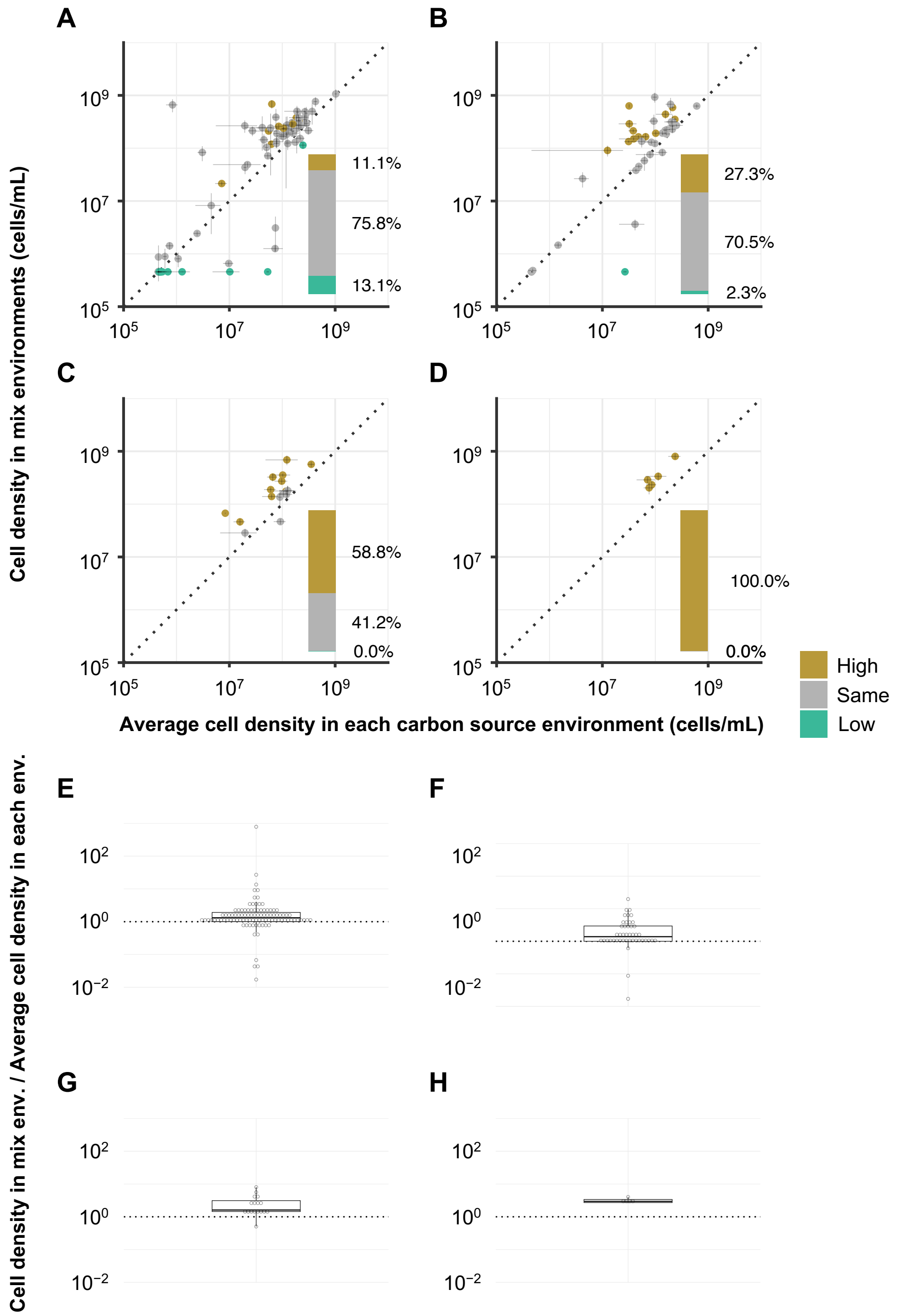


**Figure S14** Changes in cell density caused by mixing carbon sources in combinations where the provided carbon source is assumed to limit growth. (**A**–**H**) Similar to **Fig. S13**, the scatterplots (**A**–**D**) and boxplots (**E**–**H**) are shown using data on cell density for combinations expected to exhaust the available carbon sources. **A**–**D** and **E**–**H** represent the cases where the number of carbon source species after mixing was 2, 4, 8, and 16, respectively.

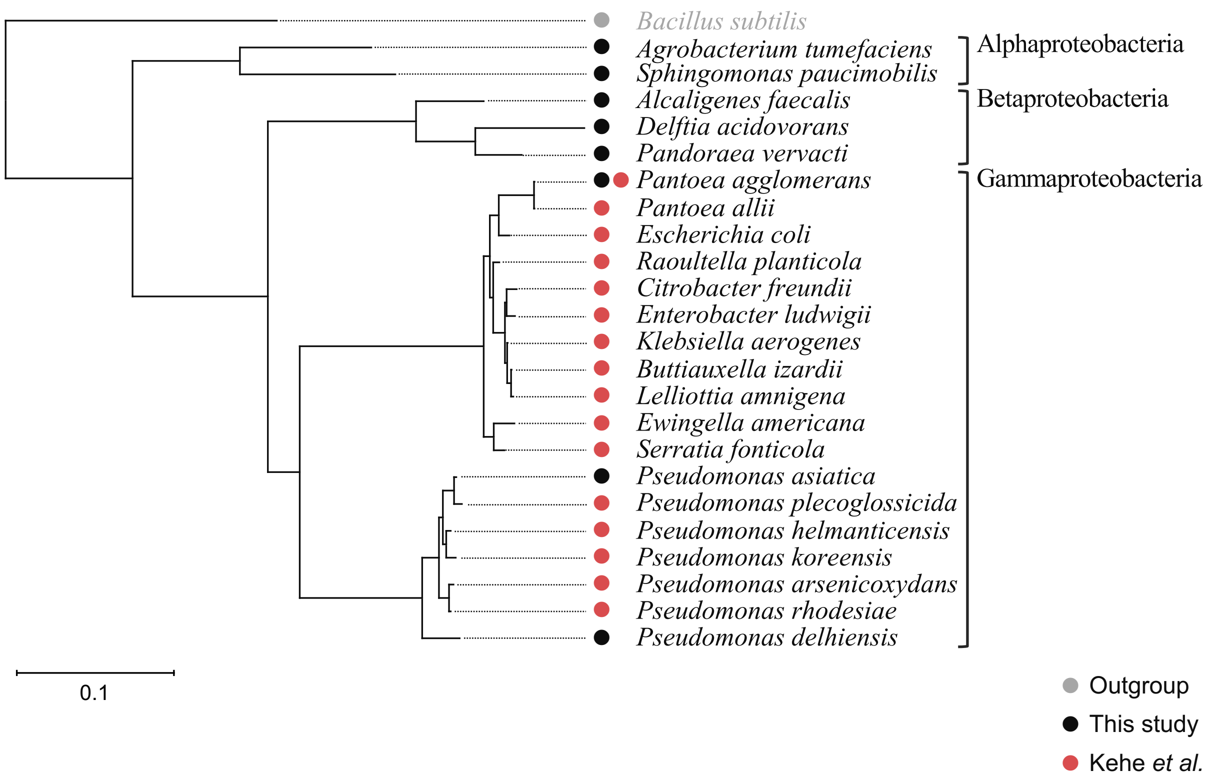


**Figure S15** Phylogenetic relationships with bacteria used in previous studies. Phylogenetic trees were created based on 16S rRNA gene sequences for both the bacteria we observed and those used by Kehe et al. (**Supplementary methods S5**). The colours of circles shown to the left of the species name refer to the description of each species.

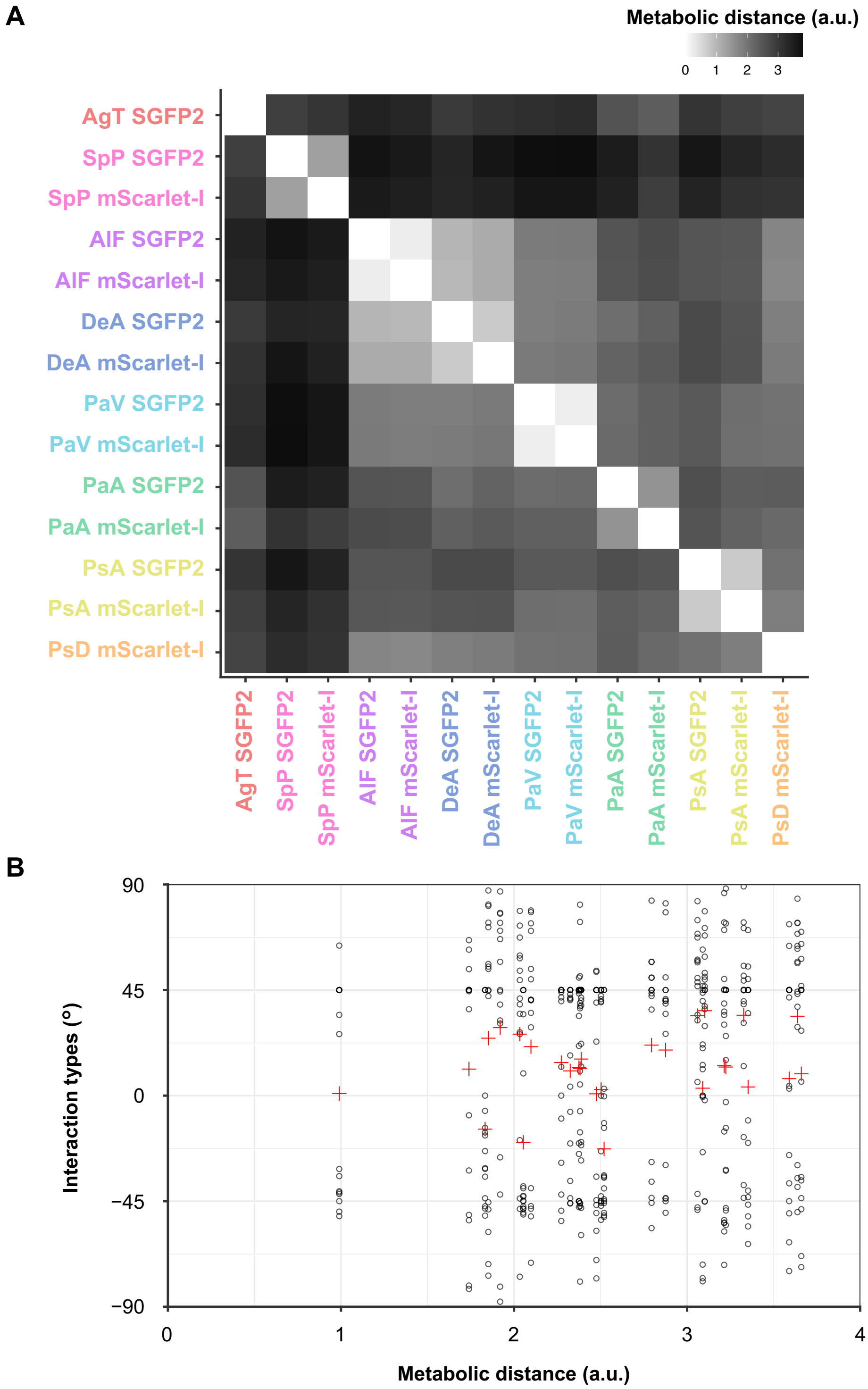


**Figure S16** Relationship between metabolic distance and interaction types. (**A**) Bacterial metabolic distance. Distance was calculated using the average cell density for each of 14 bacterial strains when mono-cultured in 32 environments. We used the cell density converted to common logarithms and calculated the ratio of each strain to its maximum growth to calculate the metabolic distance. The colours of the labels indicate the bacterial species. (**B**) Relationship between metabolic distance and the interaction types. The interaction types observed in single-carbon-source environments are indicated by round dots (Spearman's rank correlation: $\rho = 0.07,\ p = 7.13 \times 10^{-3}$). The averages of interaction types were calculated for each bacterial pair and are indicated by crossed dots (Spearman's rank correlation: $\rho = 0.16,\ p = 0.41$).

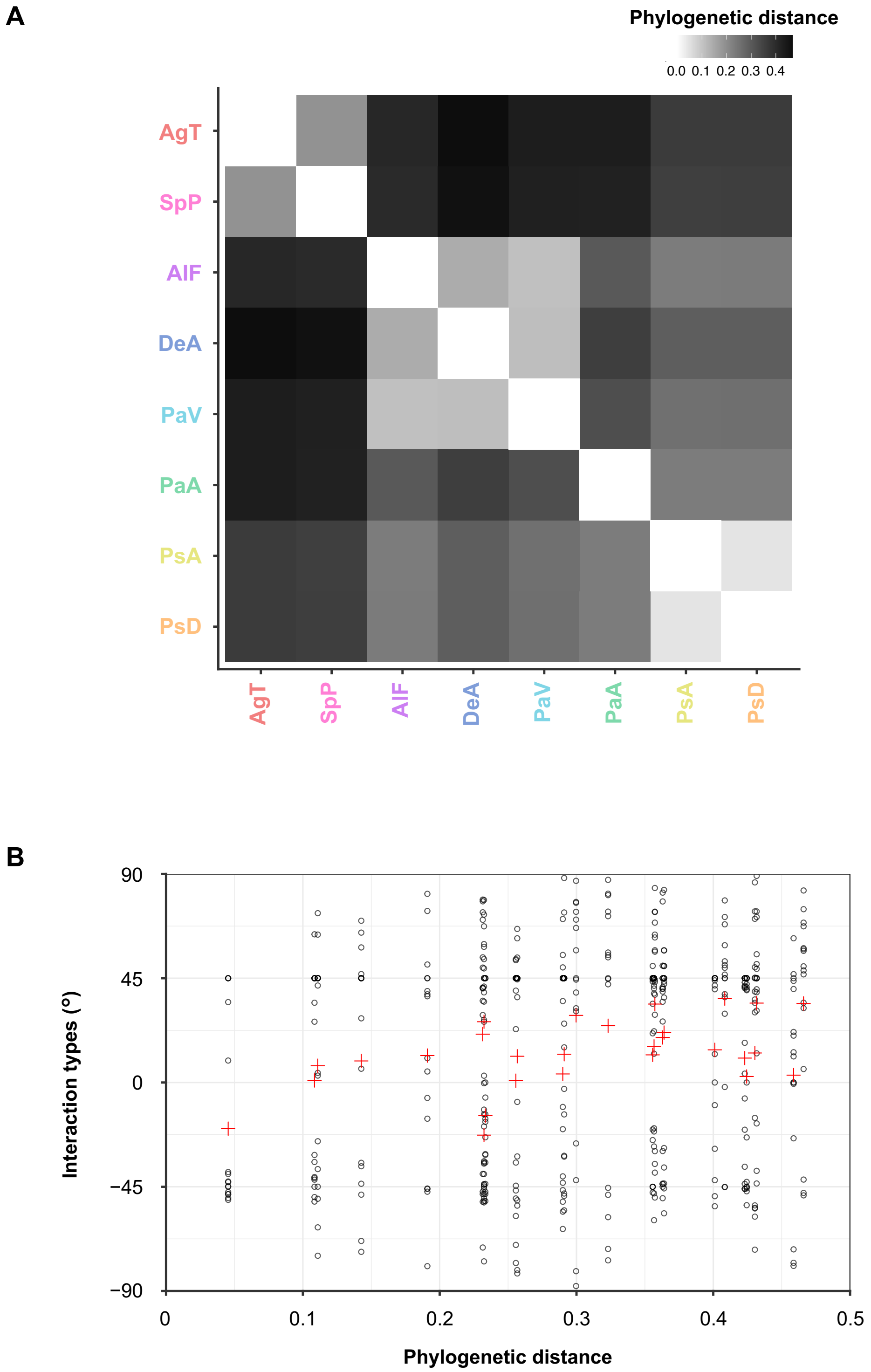


**Figure S17** Relationship between phylogenetic distance and interaction types. (**A**) Phylogenetic distance. These distance values are based on the phylogenetic tree shown in **Fig. 1A**. The colours of the labels indicate the bacterial species. (**B**) Relationship between phylogenetic distance and the interaction types. Similar to **Fig. S16**, the interaction types observed in single carbon source environments are indicated by round dots (Spearman's rank correlation: $\rho = 0.14$, $p = 3.93 \times 10^{-7}$). The averages of interaction types were calculated for each bacterial pair and are indicated by crossed dots (Spearman's rank correlation: $\rho = 0.47$, $p = 0.01$).
